# Proteomic Profiling Reveals Age-Related Changes in Transporter Proteins in the Human Blood-Brain Barrier

**DOI:** 10.1101/2024.07.26.604313

**Authors:** Xujia Zhou, Mina Azimi, Niklas Handin, Andrew Riselli, Bianca Vora, Eden Chun, Sook Wah Yee, Per Artursson, Kathleen M Giacomini

## Abstract

The Blood-Brain Barrier (BBB) is a crucial, selective barrier that regulates the entry of molecules including nutrients, environmental toxins, and therapeutic medications into the brain. This function relies heavily on brain endothelial cell proteins, particularly transporters and tight junction proteins. The BBB continues to develop postnatally, adapting its selective barrier function across different developmental phases, and alters with aging and disease. Here we present a global proteomics analysis focused on the ontogeny and aging of proteins in human brain microvessels (BMVs), predominantly composed of brain endothelial cells. Our proteomic profiling quantified 6,223 proteins and revealed possible age-related alteration in BBB permeability due to basement membrane component changes through the early developmental stage and age-dependent changes in transporter expression. Notable changes in expression levels were observed with development and age in nutrient transporters and transporters that play critical roles in drug disposition. This research 1) provides important information on the mechanisms that drive changes in the metabolic content of the brain with age and 2) enables the creation of physiologically based pharmacokinetic models for CNS drug distribution across different life stages.

## Introduction

The Blood-Brain Barrier (BBB) serves as a protective barrier that selectively regulates the entry of various substances into the Central Nervous System (CNS), including pathogens, chemical carcinogen, as well as nutrients essential for brain function and therapeutic medications. This barrier is comprised of brain microvessels (BMVs), which consists mostly of brain endothelial cells, pericytes that envelop them, and the astrocytic end-feet that form connections to these two cell layers. The basement membrane (BM), a non-cellular component synthesized by BMVs, pericytes, and astrocytes, provides structural support and facilitates cell communication [1, 2]. The tight junctions between endothelial cells, generated by multiple proteins such as claudins, occludin, junction adhesion molecules, and zonula occludens proteins, are one of the key components for proper BBB function, helping to significantly restrict the passive paracellular movement of small and large molecules [3].

To maintain brain homeostasis, the BBB possesses a variety of transport systems that enable the entry of essential molecules into the brain and prevent the accumulation of potential toxins. These systems include active efflux transport, secondary active transport, and transcytosis (receptor-mediated and absorptive-mediated transport) [4]. Active efflux transporters, mainly represented by the ATP-binding cassette (ABC) family members, P-gp (ABCB1) and BCRP (ABCG2), are highly expressed on the luminal side of endothelial cell membranes at the BBB. Although these transporters function as efflux pumps for certain endogenous metabolites, their primary role is to prevent the accumulation of xenobiotics in the brain[5]. Consequently, many CNS targeted drugs fail during drug development because of poor brain penetration due to these efflux transporters. Secondary active transporters plays an important role in bypassing the restrictive nature of the BBB, which otherwise limits the entry of macronutrients (like glucose and amino acids) and micronutrients (such as vitamins and metals)—both of which are vital for metabolism and neuronal development [6]. Secondary active transport, mainly characterized by their high selectivity, predominantly facilitate the uptake of nutrients from the blood into the brain; however, they are also capable of mediating efflux and operating as equilibrative transporters based on the concentration gradient [7]. Many of these transporters belong to the solute carrier (SLC) superfamily, including the glucose transporter SLC2A1 (GLUT1), amino acid transporter SLC7A5 (LAT1), riboflavin transporter SLC52A2, and heme transporter SLC49A2 (FLVCR2) [8–11]. These transporters are essential for normal brain development, and impairments can lead to developmental disorders, such as Glucose transporter type 1 deficiency syndrome, Brown-Vialetto-Van Laere syndrome-2, and Fowler syndrome[12].

Despite the important role of these transporters in maintaining BBB integrity and homeostasis, research exploring their variation through lifespan is limited. While previous proteomic studies have identified SLC transporters in the BBB, these studies often had small sample sizes and limited their focus on adult and elderly populations [13–15]. Evidence suggests that the BBB undergoes significant developmental changes after birth, adjusting its transport mechanisms to align with each growth stages[16]. Transcriptomic and immunohistochemical research focused on early BBB development has been explored, yet they are limited by both small cohorts and the well-documented gap between mRNA expression and protein levels and activity—the latter being the key modulators of biological functions and metabolic pathways[17–19]. Furthermore, while the abundance of transporters and BBB permeability is known to influence drug exposures in the brain, most pharmacological studies have been conducted in adult patients. In fact, despite growing evidence indicating that crucial organs involved in drug absorption, metabolism, distribution, and excretion continue to mature after birth and then decline their functions in elderly population, 50 to 90% of drugs prescribed to children or elderly are based on safety and efficacy data from clinical studies which only include adults under 65 years old, leading to a higher incidence of adverse drug reactions and therapeutic failures in these vulnerable populations [20–22]. In addition, various neurological disorders, such as stroke, epilepsy, and Alzheimer’s disease (AD), have been associated with changes in BBB function [23–25], which can either hinder drug penetration or permit excessive drug entry. Although considerable focus has been given to the efflux pump ABCB1 (P-gp) and tight-junction proteins [26–28], the effect of these disorders on the expression levels of other vital transporters, especially those involved in nutrient penetration, and other proteins important for BBB permeability is poorly understood.

This study aims to provide a comprehensive analysis of protein expression changes at the BBB across three critical age stages — development, adulthood, and old age — focusing specifically on transporters, as their expression levels can significantly impact the pharmacokinetics of drugs in the brain as well as the brain metabolome. In addition, we present some exploratory results on the BBB protein expression for a few patients with AD. Addressing these knowledge gaps in BBB protein dynamics during development and disease is crucial to understand the effects of aging during development and senescence on the brain metabolome as well as for the development of drugs for the treatment of CNS disorders.

## Results

### Experimental workflow and overview of brain microvessel (BMV) proteome

Global proteomic analysis using LC-MS/MS was conducted on BMVs isolated from thirty-four healthy brain specimens, collected from neonates to elderly individuals, as well as from three samples from AD patients (Supplemental Table 1). For the analysis, we separated the brain samples into three age categories: development (34 weeks gestation to 3 years), adult (4 to 60 years), and elderly (over 60 years). A brief overview of our study’s workflow is illustrated in Figure 1A. In total, we identified 9005 proteins across all samples (including disease samples). To account for missing data and proteins that may only be expressed in specific age groups, we only included those proteins detected in at least 70% of samples within one or more age groups for subsequent analyses; this resulted in the identification of 6,887 proteins. 6,223 proteins, which were detected with three or more razor and unique peptides, were considered quantified and subsequently used for expression analysis. We employed PANTHER [29](www.pantherdb.org/) to categorize this BMV proteome based on their protein classes. Our BMV proteome is distributed across 13 distinct protein classes, including transporters (7.54%) and cell adhesion molecules (2.16%) (Fig. 1B). Furthermore, gene ontology (GO) enrichment analysis of this BMV proteome revealed BBB-associated GO terms such as *cell substrate adhesion*, *cell junction assembly*, and *transport across the BBB* (Fig. 1C). Major tight junction proteins and integrins (e.g. CLDN5, ITGB1, TJP1) are expressed in our BMV proteome and age-dependent variation in the expression was observed for some of them. Notably, Claudin 11 (CLDN11), which has been shown to colocalize with CLDN5, exhibits significantly lower expression in Development group (Supplemental Fig. 1) [30]. An unsupervised principal component analysis (PCA) was conducted to explore the underlying variations within our dataset. We found that the BMV proteome can be effectively categorized by age groups based on the first two principal components (PCs) (Fig. 1D). The PCA loading plots allowed us to identify proteins related to the observed clustering. These proteins, including cytoskeletal-associated protein 4 (CKAP4) and C-X-C motif chemokine ligand 12 (CXCL12), are known to play crucial roles in BBB permeability (Supplemental Fig. 2A) ([31] [32]). Our BMV proteome confirmed that these proteins are also significantly correlated with age (Supplemental Fig. 2B).

**Figure 1.**
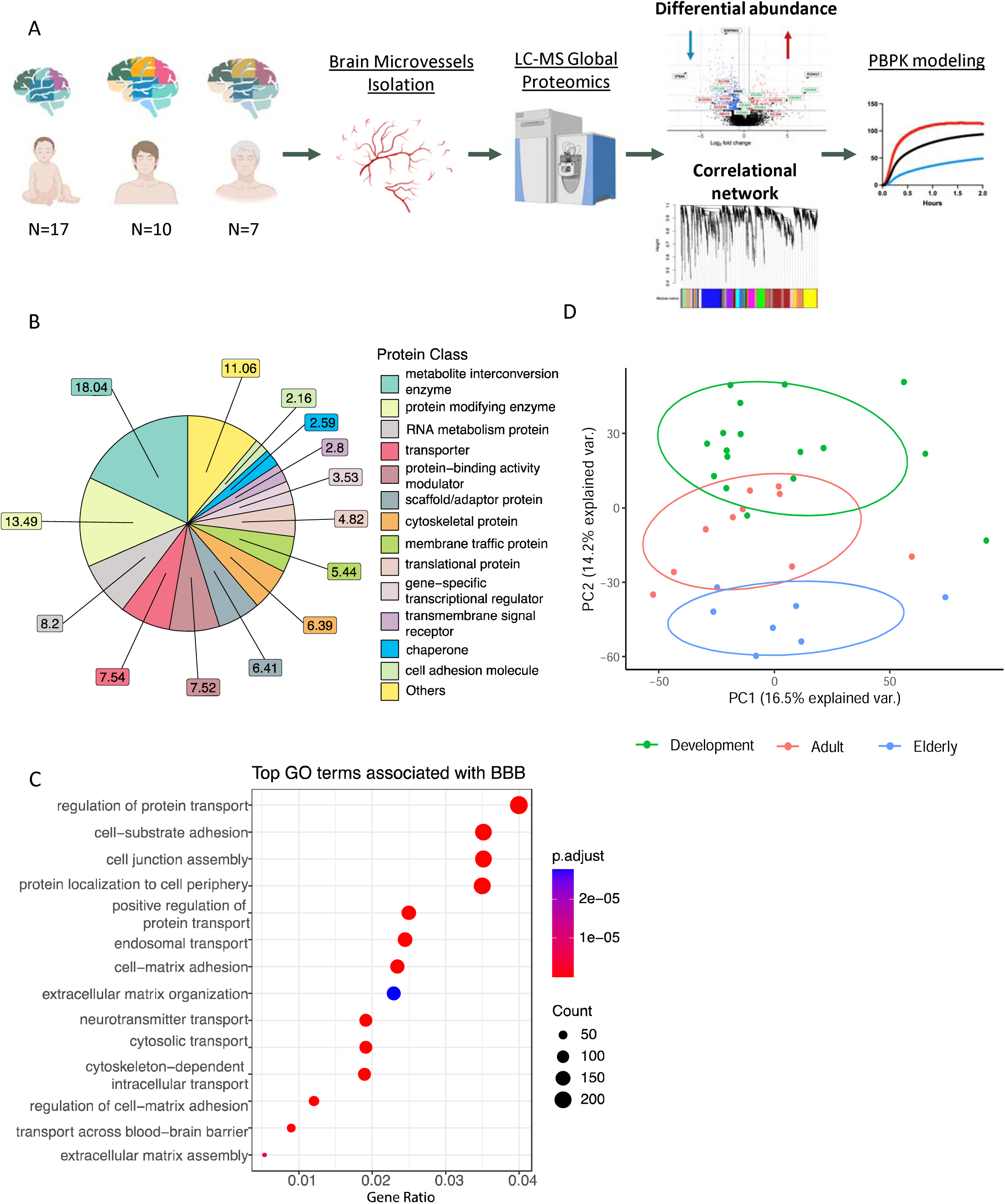
Brief experimental workflow and overview of BMV proteome. (A) Brief experimental workflow. Brain microvessels (BMVs) were isolated from frozen insular cortical bran tissue, which were then digested and analyzed using LC–MS/MS proteomic methods. Differential protein analysis and weighted correlation network analysis(WGCNA) were performed to identified proteins different between age groups and possible altered Blood brain barrier parameters were incorporated into physiologically based pharmacokinetic (PBPK) modeling simulation. (B) The PANTHER protein class of the identified proteins in BMV proteome. (C) Top BBB related GO terms associated with our BMV proteome are shown. X-axis is the gene enrichment ratio (Gene ratio) and the bubble size indicates the numbers of proteins associated a biological process GO term, with color maps the FDR value (p.adjust, q-value) of the enrichment analysis. (D)Principal component analysis (PCA) on BMV proteomic dataset showing the separation between different age groups.

### BMV proteome reveals expression patterns that are indicative of brain development stages

Differential expression was assessed using a statistical t-test analysis; more specifically, we aimed to identify proteins with significantly altered abundance levels (Absolute Log2 Fold Change > Log2(1.5), P < 0.1) between the Development and Adult groups. There was a total of 258 proteins with significantly increased abundance as individuals transition into adulthood (Fig. 2A, red circles). Gene Ontology (GO) enrichment analysis of these proteins underscored their substantial link to BBB functionality, illustrating their involvement in key biological processes like *cell substrate adhesion*, *cell matrix adhesion*, and *the regulation of metal ion transport* (Fig. 2B). COL4A1 and COL4A2, which encode the alpha1 and alpha2 chains of type IV collagen respectively, are known to be the major components of the basement membrane and play crucial roles in the stabilization and maintenance of blood-brain barrier (BBB) integrity [1]. Interestingly, while these two proteins were also the most abundant collagens in adulthood within our dataset, we found a significantly increased expression with age especially as individuals matured into adulthood (COL4A1, P-value=0.031; COL4A2, P-value=0.017) (Fig. 2A). Consequently, a notable shift was observed in the dominance of collagen families; specifically, Collagen VI (COL6A1, P-value= 0.075; COL6A3, P-value= 0.048) was identified as the predominant collagen during the developmental stages (Fig. 2C). Although the roles of COL4A3(P-value=0.026) and COL4A4(P-value= 0.0077), whose dysfunction leads to renal disease, have not been extensively explored in the brain, both proteins are significantly enriched in the Adult group (19.6-fold and 117.1-fold increase respectively) [33]. Analysis of our BMV proteome identified 29 collagens, with 37.9% (11 collagens) showing a negative correlation with age and 24.1% (7 collagens) exhibiting an increase during the developmental phase (Fig.2D). This indicates that alterations in the components of the basement membrane throughout development can potentially affect the permeability of the BBB through passive diffusion as individuals age.

**Figure 2.**
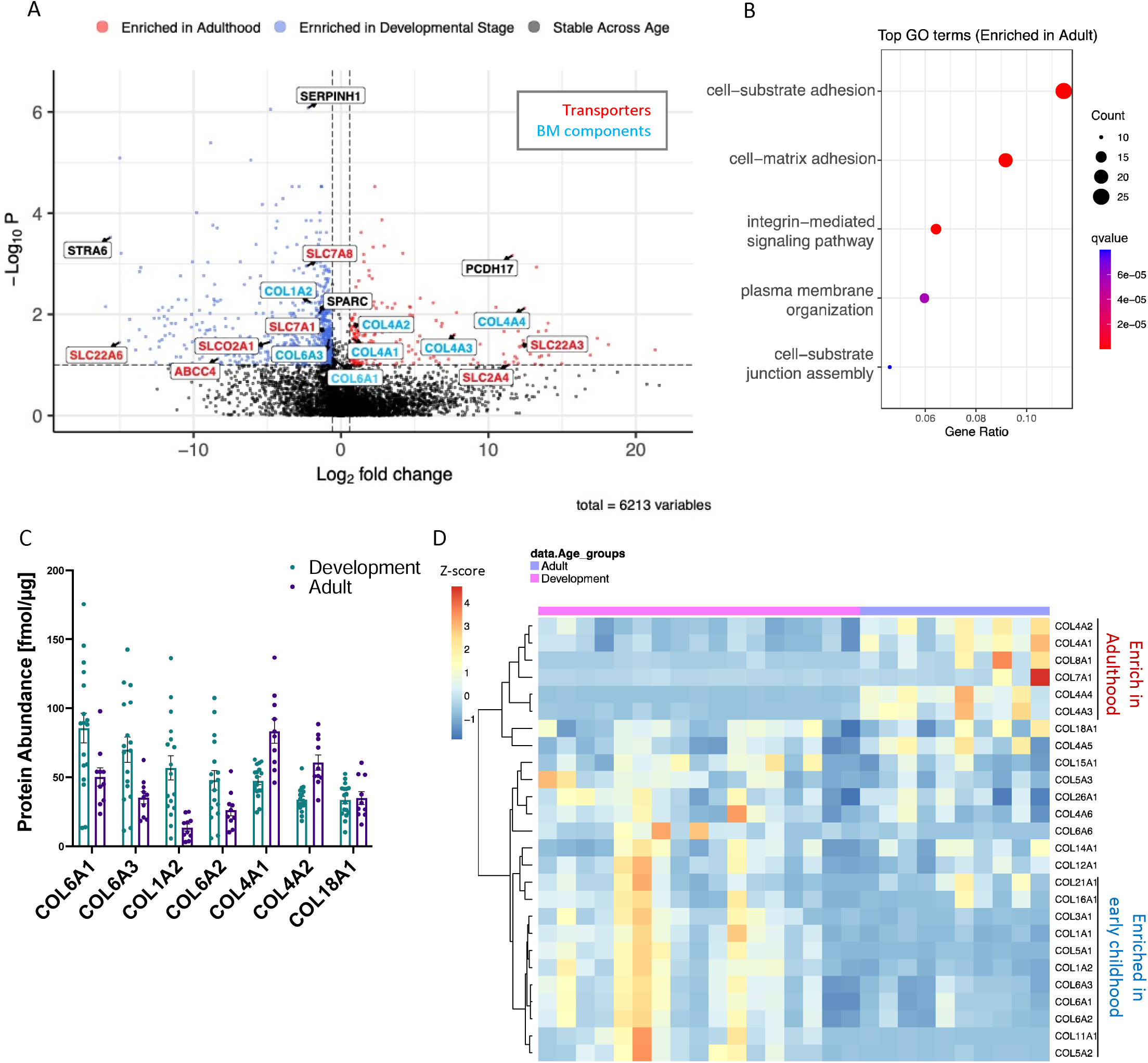
Differential expression of discovery BMV proteome through early childhood. (A) Volcano plot displaying the log2 fold change (x axis) against the t test–derived −log10 statistical P value (y axis) for all proteins differentially expressed between Development group (N=17) and Adult group (N=10) of the B proteome. Proteins with significantly decreased levels in Adults (P < 0.1) are shown on the left side, while the proteins with significantly increased levels through developement are shown on the rigMVht side. Transporters are labeled in red and proteins important for BBB integrity are labeled in green. (B) Top GO terms associated with proteins significantly increased with age are shown. X-axis is the gene enrichment ratio (Gene ratio) and the bubble size indicates the numbers of proteins associated a biological process GO term, with color maps the FDR value (p.adjust, q-value) of the enrichment analysis. (C) Bar graph showing collagens highly expressed in the BMV proteomic dataset, ordered by expression levels with the highest in the Developmental group. (D) Protein expression of Collagens which are expressed in our proteomic dataset and exhibit positive(red) or negative(blue) correlation with age. Scale represents the row Z-score (each row presents one protein), which is calculated by taking each individual’s protein expression, subtracting the mean expression, and then dividing by the standard deviation of that protein.

Conversely, there were a total of 837 proteins that were significantly enriched in developmental stage (Fig. 2A, blue circles). These included SERPINH1(a collagen-specific ER chaperone; P-value=9.18 × 10^−7^) and SPARC (an extracellular chaperone required for spatial assembly of collagen IV; P-value= 0.00980) [34]. This decrease further supports the dynamic development of the basement membrane during early childhood. Moreover, GO analysis revealed that the proteins that were decreased through development were predominantly involved in *extracellular matrix organization* and *mRNA processing* (Supplemental Fig. 3A). A similar trend has been observed when comparing human neonatal brain endothelial cells from individuals aged 15–18 and 20–23 gestational weeks [18]. Our results here support that these protein expressions continue to change during early development to adulthood, indicating BBB development is still ongoing after birth and that the metabolic process of RNA in early childhood is different from adults.

### Network BMV proteome reveals modules linked to barrier integrity and the BBB transport system

We subsequently performed a network analysis on the discovery BMV proteome using the weighted gene co-expression network analysis (WGCNA) algorithm, which organizes the dataset into modules of proteins with similar expression patterns [35]. This method of analysis is often used to discover relationships between network, gene, and sample traits in a system with higher sensitivity, capturing proteins with smaller fold changes or with lower abundance. The WGCNA identified 30 modules (M1 to M31) of co-expressed proteins ranked and numbered according to size from largest (M1, n = 792 proteins) to smallest (M31, n = 30 proteins) (Fig. 3A). The biology represented by each module was determined using network analysis provided by Cytoscape [36] (https://cytoscape.org/, Supplemental Table 6). To assess whether a module was related to the age shift, we correlated each module eigenprotein, or the first principal component (PC) of module protein expression, to each sample’s specific age (in days). We observed that 10 modules show a significant positive correlation (> 0.1) and 7 modules show a significant negative correlation (< -0.1) through the early development to elderly stages. Modules which positively correlated with age consisted of protein groups related to protein maturation and positive regulation of endopeptidase activity. Modules which negatively correlated with age include protein groups related to regulation of mRNA processing and posttranslational protein targeting to endoplasmic reticulum membrane. Since our BMV proteome includes samples from a large range of ages, and many proteins do not have a linear change throughout a person’s lifetime, in order to further dissect how these modules are altered through development and in the aging process respectively, we compared the module eigenprotein across age groups (Development, Adult, and Elderly). The module linked to collagen fibril organization (M7) which includes basement membrane components such us COL1A2 (Collagen type I alpha 2 chain), COL11A1 (Collagen type XI alpha 1 chain), and LUM (Lumican), shows significantly enriched expression in the Development group (Fig. 3B). This is consistent with the GO enrichment analysis we did utilizing the differentially expressed proteins among the Development and Adult groups. While the selective barrier function of the BBB is derived from the tight junctions between endothelial cells of the BMVs and the basement membrane surrounding, transporters are another crucial component for restricting entry of many xenobiotics into the brain, as well as for delivering essential nutrients for proper brain development and homeostasis. Our analysis identified a significant alteration in the module linked to transporters across the BBB (M12) during early developmental stages, with nutrient transporters such as the amino acid transporters, SLC7A1 (CAT1), SLC7A5 (LAT1), and the docosahexaenoic acid transporter, SLC59A1(MFSD2A), exhibiting age-dependent changes (Supplemental Table 2).

**Figure 3.**
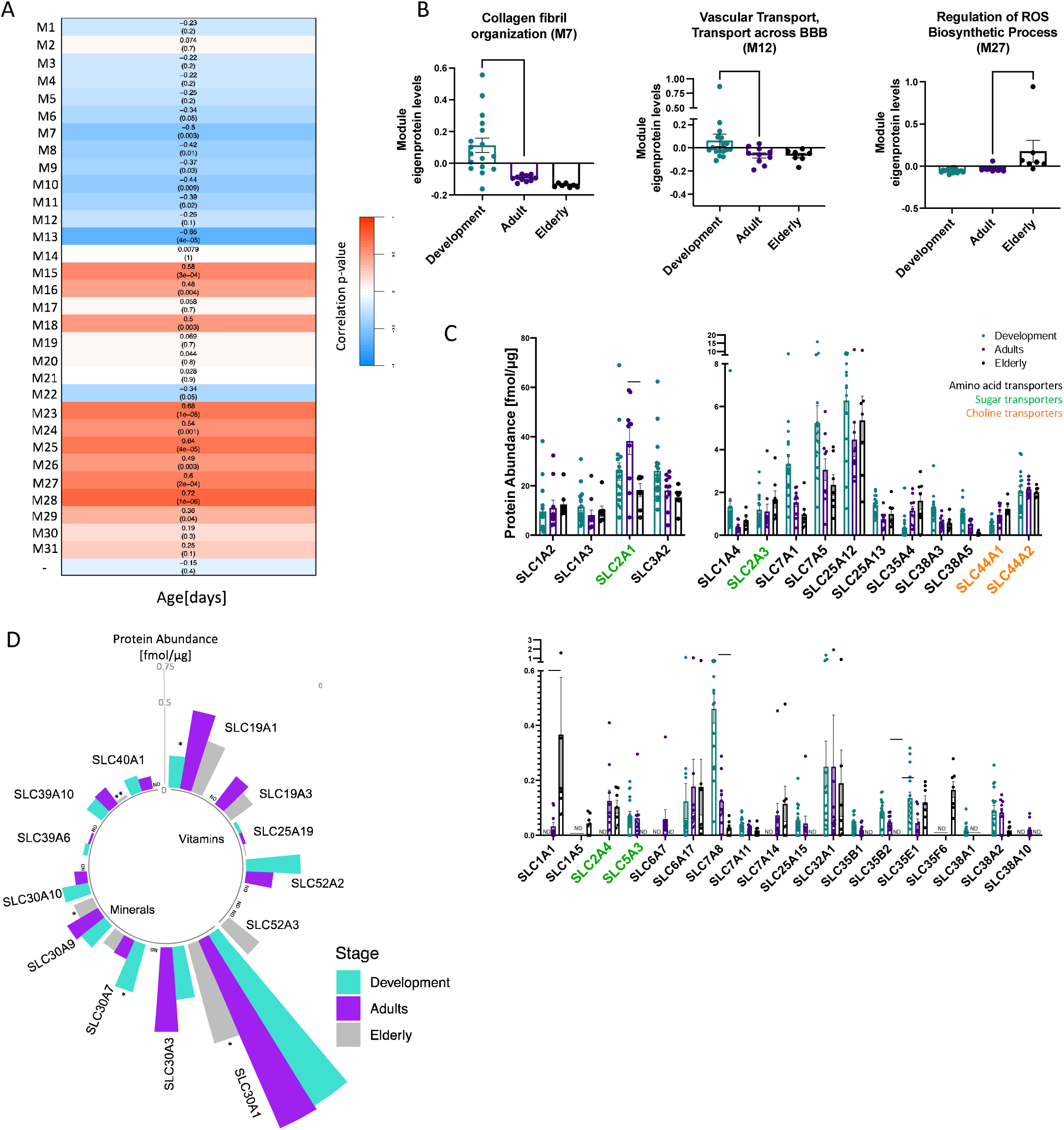
BMV proteins co-expression network identified age-dependent modules which are important for BBB function. (A) The correlation between modules and age. Heatmaps shows the correlation between eigengene and age and each cell contains the corresponding correlation followed by p-value. (B) Module eigenprotein levels by age groups (Development, Adult, Elderly) for the three blood brain barrier related modules. Modules are grouped by different age groups. (C) Macronutrient transporters in our BMV proteomic dataset are shown in bar graph. Amino acid transporters are labeled as black, sugar transporters are labeled as green and choline transporters are labeled as orange. (D) Micronutrient transporters in our BMV proteomic dataset are shown in circular bar graph, bars represented the mean of the transporter expressions in different age groups. For the bar graphs Kruskal–Wallis tests followed by Dunn’s post hoc test were used to compare the mean of each age groups with the mean of the Adult group. Data are represented as mean ± SEM and each points represent one sample. *P < 0.05, **P < 0.01, ***P < 0.001. ND, not detected in more than 30% of samples in specific age group.

### Nutrient transporter expression in the BBB shows age dependence

The human brain goes through significant changes postnatally. While a newborn already has most of its lifelong neurons, the brain doubles in size the first year due to an accelerated rate of myelination and the formation of synapses during early childhood ([37], [38]). Proper nutrition and availability of nutrients to the CNS are crucial for normal brain and neurocognitive development. We identified a total of 56 SLCs involved in nutrient transport in the human BMV proteome. Twenty-five of the 56 have been quantified for the first time through a global proteomic analysis of the human BMVs[15, 39]. Moreover, our proteomic analysis revealed that the BBB, which controls the delivery of essential nutrients to support brain and neuron development, has differential protein expression during early development, including that of multiple nutrient transporters. Amino acids such as l-arginine and l-serine are required for regulating synapse formation/patterning and neurotransmitter synthesis [40]. In our dataset, we found that l-arginine transporter SLC7A1 and glutamate/l-serine SLC38A5 (SNAT5), were highly expressed in early childhood, with expression levels decreasing in adulthood (Fig. 3C). Additionally, although the group-wise comparisons did not reveal significant differences, SLC3A2 (CD98hc/4F2hc) and SLC7A5, the two most highly expressed amino acid transporters in our BMV proteome, exhibit a significant negative correlation with age (Supplemental Fig. 4).

Micronutrients such as vitamins and minerals are crucial for normal neurodevelopment. For example, riboflavin is important for neurodevelopment and deficiency in riboflavin has been associated with autism spectrum disorders (ASD) and developmental delays [41, 42]. In the elderly population, riboflavin supplement has been linked to better cognitive performance by reducing oxidative stress in the brain [43, 44]. The riboflavin transporter, SLC52A2 (Km=0.33 µM) was only detected in Development and Adult groups while another riboflavin transporter, SLC52A3 (Km=0.98 µM), which is in the same family but has a slightly higher Michaelis–Menten constant (Km), only expressed Elderly group (Fig. 3D) [38, 45]. These data suggest that riboflavin delivery to the brain may change depending on age, with the increased delivery occurring during development. Interestingly, while zinc transporters are among the most highly expressed micronutrient transporters in BMV throughout life, SLC30A1, the most abundant zinc transporter, shows significantly lower expression in the elderly population. Recently identified as the primary transporter for choline uptake into the brain, SLC49A2 has been found in BMVs in both Development and Adult groups but was undetected in the Elderly group (Supplemental Table 2)[46]. This aligns with clinical observations of reduced choline uptake into the brain in the elderly population[47]. These observations suggest that delivery of specific macro- and micro-nutrients to the brain changes with maturation and aging.

### Transporters linked to Mendelian Diseases with neurological manifestations are predominantly expressed in early childhood

Given the pivotal physiological roles of SLC transporters in maintaining brain homeostasis, and their critical involvement in delivering essential nutrients for brain development, it is not surprising that a malfunction in a single SLC transporter can lead to severe neurological disorders. Mutations in 20% of known SLC transporters in humans are associated with Mendelian diseases, with 39 of which were identified in our BMV proteome ([48], Table 1). Especially, SLC transporters associated with mendelian diseases that present neurological symptoms whose primary substrates are crucial for brain development are prominently expressed in early childhood. Mutations in the SLC1A4 transporter lead to spastic tetraplegia, a thin corpus callosum, and progressive microcephaly, an neurodevelopmental disorder with severely impaired global development in early infancy due to disrupted serine transport into the brain. We identified that SLC1A4 is significantly enriched in the Development group (Fig. 3C, Table 1, Dunn’s multiple comparison test p-value (Dunn’s p-value) =0.042) and decrease 70% of its expression as we are getting into adulthood. Thyroid hormones also play a critical role in brain development, impacting brain function throughout life. Mutations in the thyroid hormone transporter, SLC16A2, are linked to Allan-Herndon-Dudley syndrome (AHDS), a disease characterized by severe intellectual impairment, dysarthria, athetoid movements, muscle hypoplasia, and spastic paraplegia [49]. We discovered that the expression of SLC16A2 in BMVs was highest in the very young (Dunn’s p-value =0.0041) and exhibited a sequential reduction with age. (Table 1). These reemphasize that the BBB adjusts the transport of macronutrients into the brain in response to developmental needs. Moreover, manganese (Mn) is an essential trace nutrient for neuron function, however, excess Mn in the brain leads to neurotoxicity [50]. Mutations in either a cell surface–localized manganese efflux transporter SLC30A10 or in a manganese influx transporter SLC39A8, result in mendelian diseases with neurological symptoms ([51], [52]). We quantified that the expression of SLC30A10 on BMVs was significantly enriched in the Development group (Dunn’s p-value =0.012) while SLC39A8 was highest in adulthood (Fig. 3C, Supplemental Table 3). Interestingly both expressions vanished in the elderly. These different expression patterns suggests that manganese concentration is rigorously regulated by transporters on BBB.

**Table 1.**
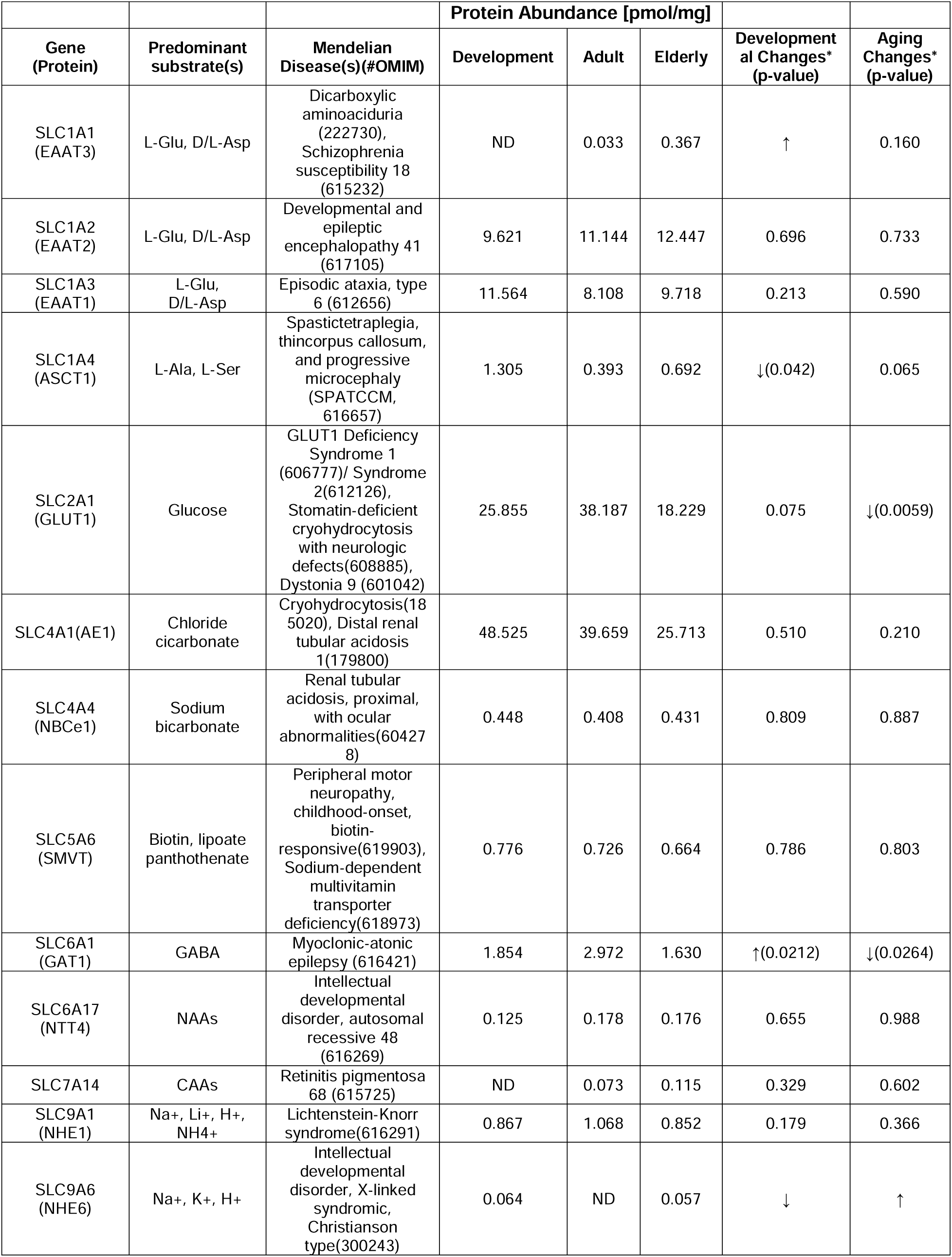
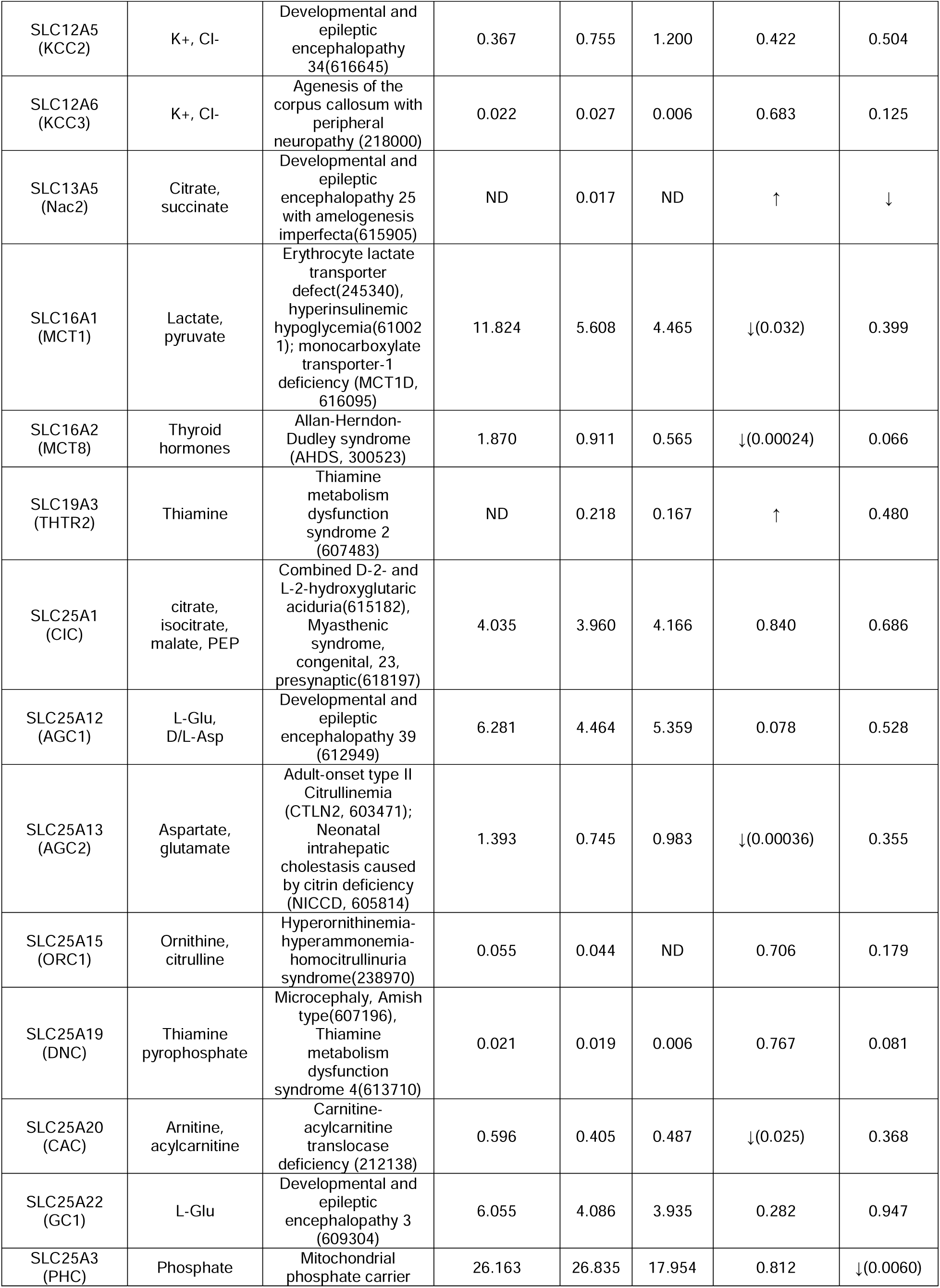
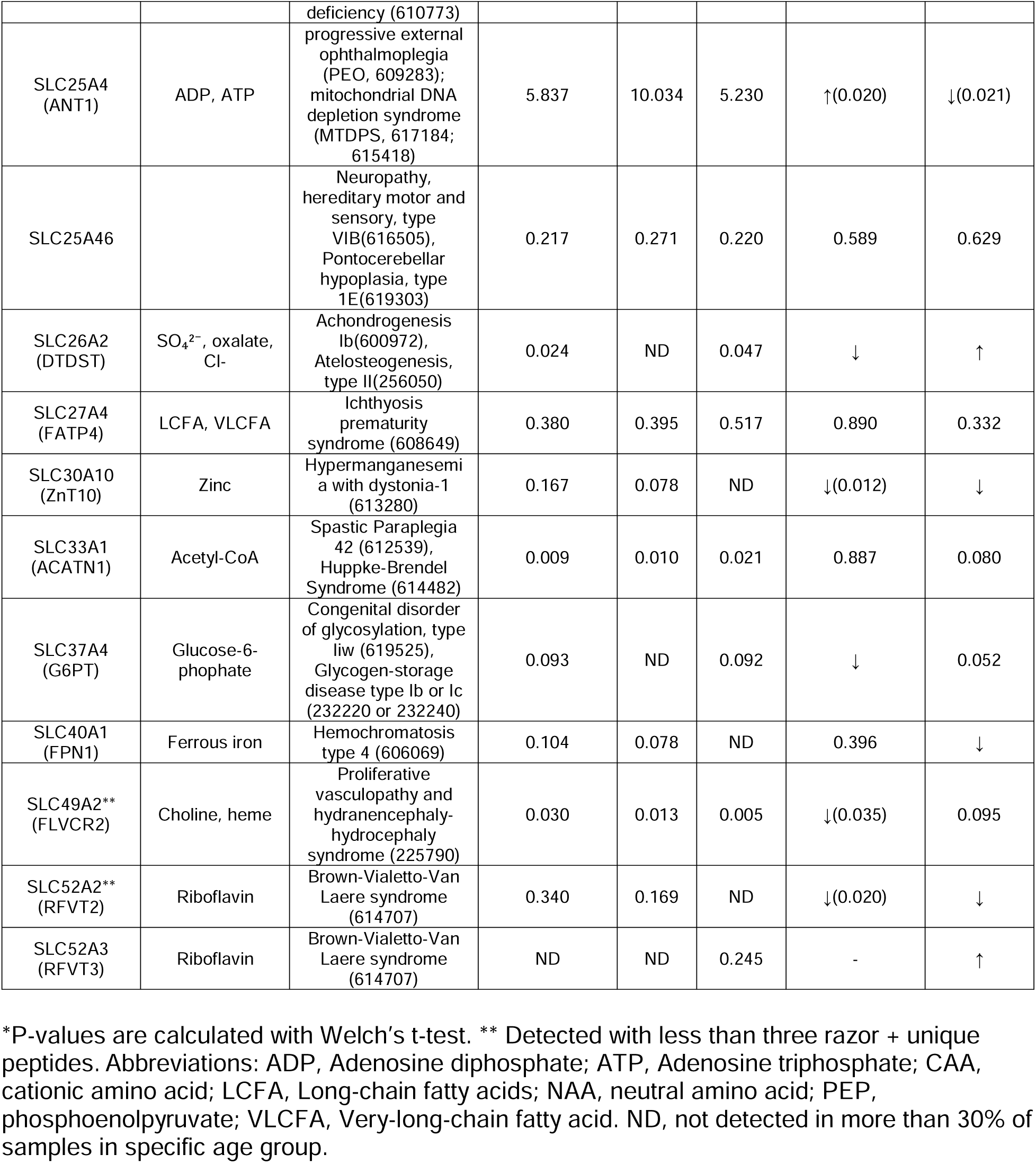
SLC transporters on BBB associated Mendelian Disease

### ADME transporters are expressed on BBB

Due to limited transport across the BBB, drugs approved to treat neurological diseases are typically restricted to lipid-soluble small molecules that can pass through the BBB via passive diffusion to reach their target in the brain. However, these drugs must also avoid efflux from transporters, such as ABCB1 and ABCG2, to maintain their brain concentrations. Drugs that initially cannot cross the BBB can be modified to be transported via inherent BBB mechanisms, such as carrier-mediated transport and receptor-mediated transport systems. SLC7A5 and TFRC which are well-known for their roles in endogenous BBB carrier-mediated and receptor-mediated drug transport respectively, both exhibited a negative correlation with age. Their expression levels were highest in the Development group and decreased with age, showing reductions of 55.2% for SLC7A5 and 78.5% for TFRC compared to the Adult group (Supplemental Table 2, 3).

ABCB1 (P-gp), ABCG2 (BCRP), ABCC4 (MRP4), SLCO2B1 (OATP2B1), SLCO1A2 (OATP1A2), and SLC29A1 (ENT1) are clinically important transporters involved in the absorption, distribution, metabolism, and excretion (ADME) of drugs within the body [53]. These transporters, known to be expressed in the BBB [54], were all identified in our BMV proteome (Fig. 4A). Consistent with prior studies, ABCB1 and ABCG2 were the most highly expressed among the ABC transporters, although no notable age-related expression differences were observed. Age-dependent expression was evident in some transporters; ABCC4 is prevalent in Development and Elderly groups, while SLCO1A2 declined with age (Fig. 4A). SLCO1A2 transport statins ([55, 56]), which are among the most commonly prescribed medications worldwide and are used in children with heterozygous familial hypercholesterolemia from as early as 8 years old ([57, 58]).

**Figure 4.**
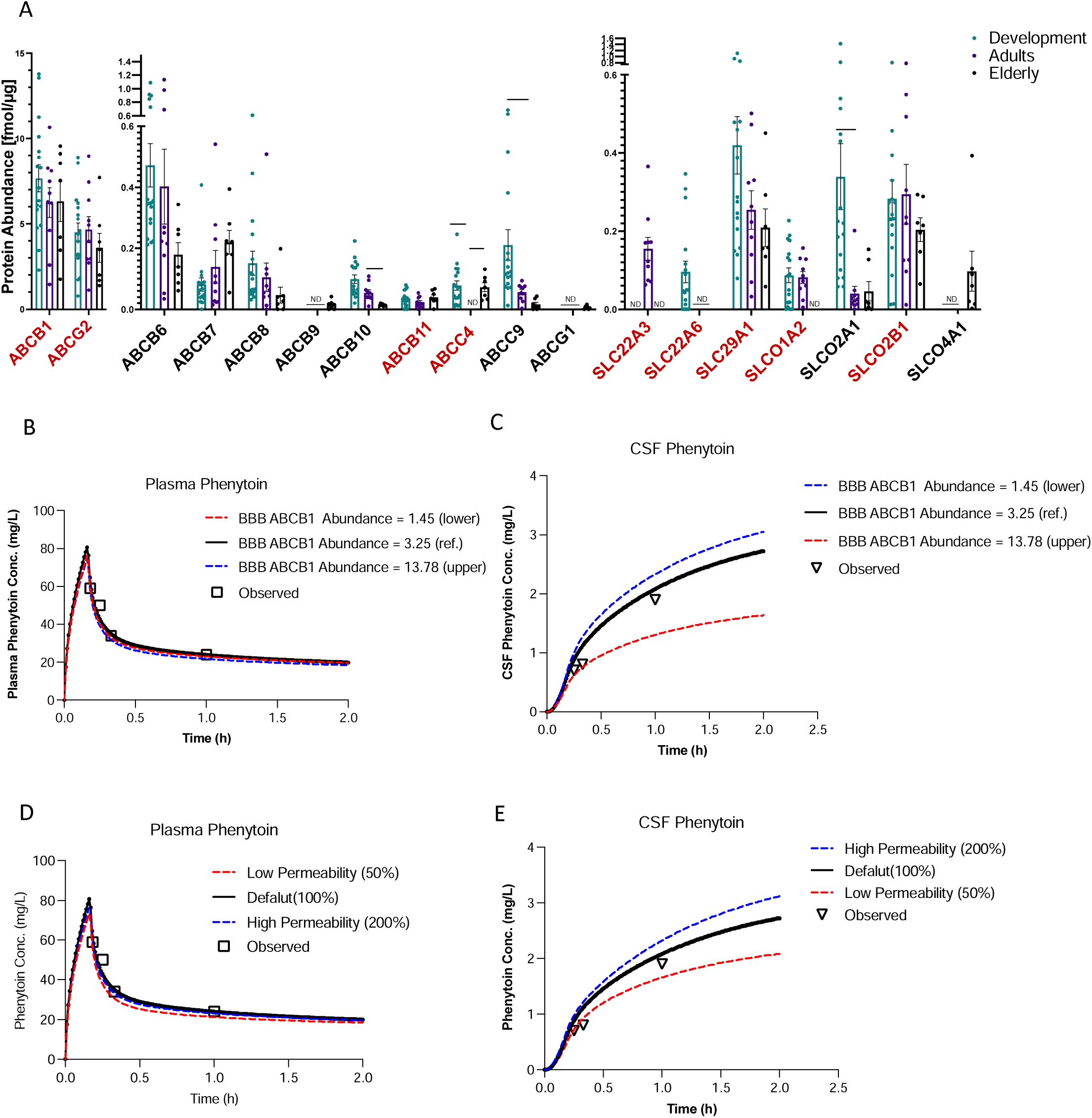
Changes in clinically important ADME transporters and BBB permeability potentially lead to different drug distribution in brain. (A) Clinically important uptake and efflux transporters (labeled red) and their family members in our BMV proteomic dataset are shown in bar graph. Kruskal–Wallis tests followed by Dunn’s post hoc test were used to assess difference between age groups in bar graph. Data are represented as mean ± SEM and each points represent one sample. *P < 0.05, **P < 0.01, ***P < 0.001. ND, not detected in more than 30% of samples in specific age group. (B, C) Phenytoin time-concentration profile in plasma and CSF with varying levels of P-gp (ABCB1) expression at the BBB. The default value is represented by the black solid line, the minimum value from BMV proteome is shown by the blue dashed line, and the maximum value from BMV proteome is indicated by the red dashed line. (D, E) Phenytoin time-concentration profile in plasma and spinal CSF with different BBB permeability. The default value is shown by the black solid line, 200% of the default BBB permeability is represented by the blue dashed line, and 50% of the default BBB permeability is shown by the red dashed line.

Other transporters that play a role in ADME in the BBB also exhibited differential expression with aging. SLC22A3 (OCT3) showed higher expression levels in Adult group but not in Development and Elderly groups. On the other hand, SLC22A6 (OAT1) is expressed only in early childhood (Fig. 4A). Methotrexate (MTX) is a well-known substrate of SLC22A6, and studies have shown that ABCC4 also plays an important role in the brain distribution of MTX [59, 60]. MTX is one of the most commonly used chemotherapy drugs for acute lymphoblastic leukemia (ALL), the most frequently diagnosed cancer in patients under 15 years of age in the United States. However, MTX is associated with significant drug-induced neurotoxicity, as well as long-term neurological deficits. Younger patients in comparison to older age patients are at reduced risk for MTX-induced neurotoxicity [61, 62]. Age-dependent changes in transporters for MTX including OAT1, ABCC4, ABCB1 and ABCG2 may account for differences in risk for MTX-induced neurological toxicities with age. Interestingly, SLC22A8 (OAT3), which is highly expressed in the rodent BBB [63], was undetectable in our samples, highlighting species differences in BBB transport proteins. Supplemental Table 4 provides a list of ABC and SLC transporters that play important roles in drug disposition, but do not appear to be expressed to a detectable degree in our BMV proteome.

### Simulations demonstrate the impact of transporter expression and BBB permeability on CSF drug exposure

As proof of concept, we employed physiologically based pharmacokinetic (PBPK) modeling and simulation with a published four-compartment, permeability-limited brain model to explore how individual differences in BBB passive permeability and ABCB1 expression affect distribution of the anti-epileptic drug phenytoin to the CNS [64]. The simulations incorporated data on ABCB1 expression obtained from our proteomic analysis, as well as BBB permeability, to demonstrate the effects of variability in these parameters. The simulation results, depicted in Figure 4 B-E, demonstrate the concentration-time profiles of phenytoin in plasma and cerebrospinal fluid (CSF). Notably, our simulations show that changes in ABCB1 expression and BBB permeability lead to pronounced changes in phenytoin exposure within the spinal CSF, with minimal impact on plasma phenytoin levels. Specifically, reduced ABCB1 expression at the BBB (by 2.24 fold) results in a predicted increase in the CSF exposure of phenytoin (default 3.25 vs. lowest value in our dataset 1.45) and in particular, a 12.2% predicted increase in the area under the curve (AUC) in the CSF (Fig. 4C, blue line). Conversely, a 4.24-fold increase in ABCB1 expression (default 3.25 vs. highest value in our BMV proteome 13.78), results in a predicted 37.5% reduction in CSF phenytoin AUC (Fig. 4C, red line). Additionally, lower BBB permeability results in a predicted decrease in phenytoin exposure in the CSF, for example, a two-fold reduction in BBB permeability results in a reduction in the CSF AUC by 21.0% (Fig. 4E, red line). Conversely, higher BBB permeability results in a 12.3% increase in phenytoin CSF AUC (Fig. 4E, blue line). Given the narrow therapeutic range of phenytoin, elevated CNS concentrations can lead to neurotoxicity, manifesting as symptoms ranging from mild nystagmus to ataxia, lethargy, and in severe cases, coma and death [65]. Interestingly, because of saturable metabolism, phenytoin has been associated with high interindividual differences in plasma levels [66]. Here we find that the high variability in the expression levels of ABCB1 in BMVs from different individuals may also contribute to the observed large interindividual differences in both efficacy and toxicity of phenytoin. Conventionally, serum phenytoin levels are monitored to ensure appropriate dosing and minimize adverse effects. However, our study suggests that serum phenytoin concentrations may not accurately indicate the risk for neurotoxicity, as they do not adequately reflect brain drug levels, which are attributable to differences in BBB permeability and transporter activity.

### Proteins associated with higher risk of AD are correlated with age and differentially expressed in AD patients

AD presents a significant challenge in healthcare due to the lack of effective curative treatments. Among the various risk factors for AD, aging is the most significant contributor. Differential protein expression between the Adult (N=10) and Elderly groups (N=7) was analyzed using the same methodology previously described. We identified a total of 168 proteins with significantly increased abundance and 427 proteins with significantly decreased abundance during the aging process (Fig. 5A). These differentially expressed proteins include several well-known proteins which are known to play important roles in the aging process, such as Apolipoprotein D (APOD), a secreted lipocalin important for lipid metabolism that has been known to increase in aging human brains [67]. LRRC32, a protein involved in the regulation of transforming growth factor beta and whose increased expression has been reported to correlate with cognition impairment and inflammation response in the brain, also showed differential expression in protein levels in Elderly group[68]. GO enrichment analysis of these 168 proteins with significantly increased expression revealed strong links to *cell substrate adhesion* and *leukocyte migration* (Supplemental Fig. 3B). Conversely, the 427 proteins with significantly decreased expression were strongly associated with *endocytosis*, *cell junction maintenance*, and *zinc ion transmembrane transport* (Fig. 5B). These protein expression shifts indicate the permeability and function of BBB may be broken down with aging and may also lead to a higher risk for the onset of neurodegenerative diseases [69, 70].

**Figure 5.**
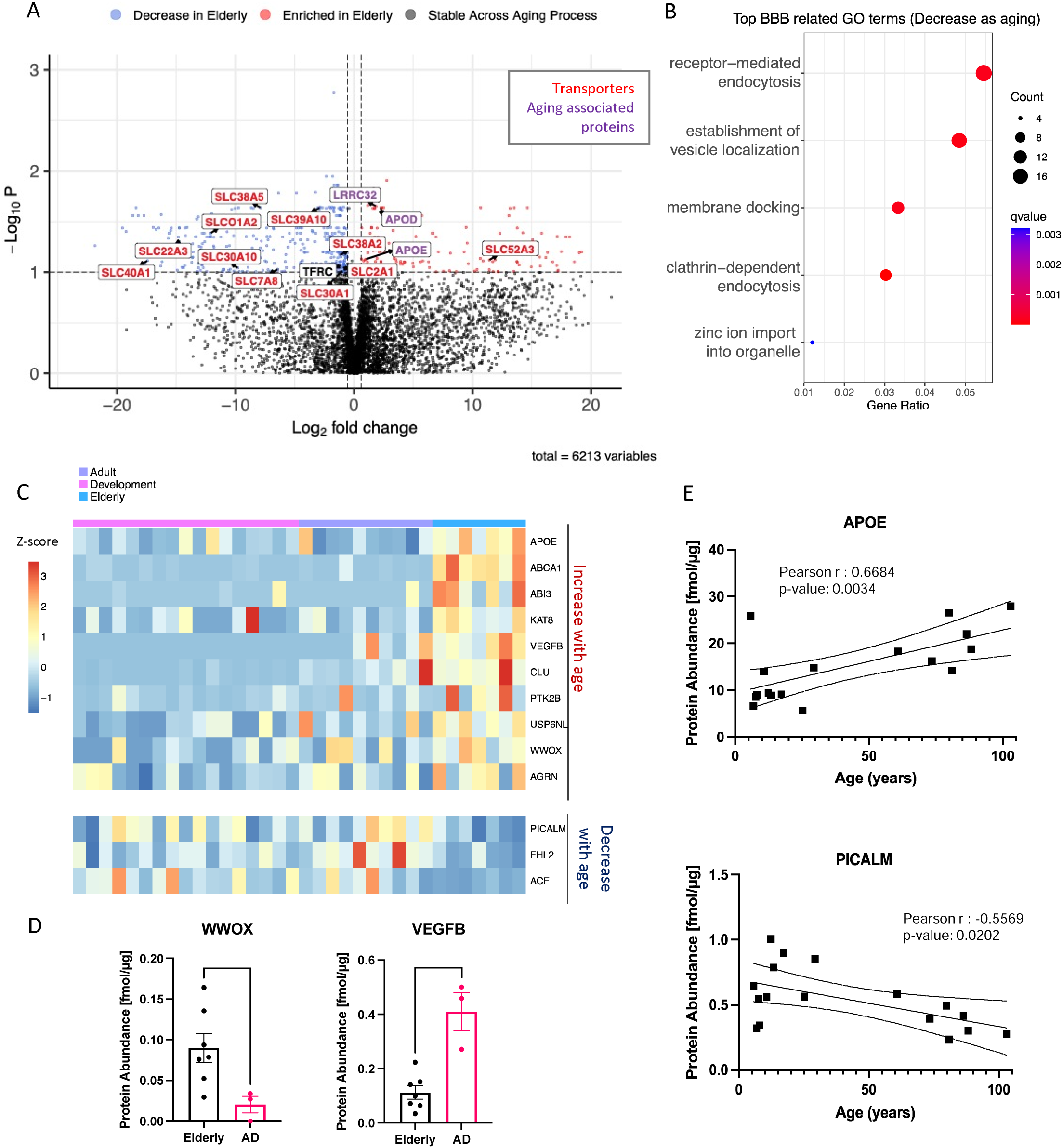
Proteins Differentially Expressed During Aging Tend to Be Associated with Alzheimer’s Disease. (A) Volcano plot displaying all proteins differentially expressed between Adult group (N=10) and Elderly group (N=7) of the BMV proteome. Proteins with significantly decreased levels in elderly population (P < 0.1) are labeled in blue, while the proteins with significantly increased levels with aging are shown in red. Transporters are labeled in red and proteins known to play crucial roles in aging are labeled in purple. (B)Top BBB related GO terms associated with proteins significantly decreased with age are shown. X-axis is the gene enrichment ratio (Generatio) and the bubble size indicates the numbers of proteins associated a biological process GO term, with color maps the FDR value (p.adjust, q-value) of the enrichment analysis. (C) Protein expression of AD GWAS genes which are expressed in our proteomic dataset and exhibit positive(red) or negative(blue) correlation with age. Scale represents the row Z-score (each row presents one protein), which is calculated by taking each individual’s protein expression, subtracting the mean expression, and then dividing by the standard deviation of that protein.(D) Protein abundance levels of WWOX and in healthy elderly population or in patients with AD. Expression difference between disease condition were assessed by student T test. *P < 0.05, **P < 0.01, ***P < 0.001 (E) Protein abundance of APOE and PICALM through aging are described as simple linear regression model. Individual curve are presented in (solid lines). Dashed lines represent the 95% confidence bands.

To explore whether proteins that change expression during aging might be linked to neurodegenerative diseases, we examined 45 AD-related genes identified from Genome-Wide Association Studies (GWAS) [19] within our BMV dataset. Of these, 22 genes were detected in our dataset, and 12 displayed age-dependent expression patterns, highlighting their potential roles in AD pathology (Fig. 5C, Supplemental Fig. 5). WW domain-containing oxidoreductase (WWOX), recognized as an AD risk factor, plays a pivotal role in the disease’s neurodegenerative processes. Our study indicates a reduction in WWOX expression in the AD patient compared to the healthy, elderly controls [71](Fig. 5D). Furthermore, aging is acknowledged as a principal risk factor for AD. Pathologically, senescent cells accumulate proteins with age; however, the aging process of BMVs is still largely unknown, particularly with regards to its link to AD risk. In our BMV dataset, six AD GWAS genes demonstrate age-specific correlations, underscoring their potential contribution in the role of aging in AD. Among these genes, Apolipoprotein E (APOE) is notable for its role in lipid metabolism and significant contribution to AD pathology [72]. Its expression, known to increase with age, is associated with key AD features such as amyloid-beta aggregation and clearance [73]. Consistent with previous studies, in our proteomic dataset, APOE shows high expression in BMVs, with a positive correlation with age (Fig. 5E). Additionally, PICALM, one of the most significant loci identified in AD susceptibility GWAS, encodes the phosphatidylinositol binding clathrin assembly protein, which is involved in endocytosis and autophagy. Our data reveal a negative correlation of PICALM expression with aging (Fig. 5E). Furthermore, vascular endothelial growth factor B (VEGFB) is a crucial protein for blood-brain barrier repair and has been known to increase in expression when the BBB is damaged. Previous studies have shown a negative correlation between brain VEGFB expression and cognition scores in AD patients [74]. We observed that VEGFB was initially absent in the BBB in the Development group but significantly increased with age, with even higher levels in AD patients, consistent with BBB disruption in AD patients (Fig. 5D). These findings illuminate the dynamic expression patterns of these genes in relation to aging and AD, providing new pathways to understand the molecular drivers behind AD progression.

## Discussion

The goal of this study was to determine the developmental and aging effects on the BBB, with a focus on the expression levels of key transporters that play a role in CNS drug disposition and micro- and macro-nutrient availability to the CNS. Our key findings are:

(a) Alterations in basement membrane components and increased expression of cell adhesion molecules occur during early development; (b) Several nutrient transporters that are important for brain development are enriched during early development; (c) Certain drug transporters that are expressed on BBB exhibit age-dependent expression patterns; (d) Exploratory studies suggest that some proteins associated with AD exhibit altered expression levels in BMVs from AD patients. Each of these findings is discussed below followed by the limitations of the study.

### Alterations in basement membrane components and increased expression of cell adhesion molecules occur during early development

Unlike the cellular components of the BBB, the basement membrane has only been recently recognized for its importance in maintaining BBB permeability [1]. Age-related changes were evident in the BBB’s basement membrane. Our BBB proteome study, for the first time, comprehensively identified the collagen families composing the basement membrane across various age groups (from neonates to the elderly). Our study revealed that the predominant collagen families in the basement membrane differ between the early developmental stage and adulthood. In early childhood, the Collagen VI family (COL6A1 and COL6A3) is the most highly expressed, whereas in adulthood, the Collagen IV family (COL4A1 and COL4A2) is predominant. These 2 collagens are distinct in terms of their morphology. Collagen VI is an unusual collagen comprising both collagenous and non-collagenous domains that assemble into beaded filaments, whereas Collagen IV forms a protomer fiber made of three α chains. Collagen IV is well-known to form the backbone of the basement membrane [75]. The Collagen VI family is highly expressed in various human tumors and influences cell proliferation and angiogenesis [76]. Since it is highly expressed in neonates, this aligns with the need for postnatal vascularization at the early developmental stage.

Cell adhesion molecules (CAMs) in the BBB are proteins located on the cell surface involved in the binding with other cells or with the extracellular matrix. Our study revealed that while the most extensively studied junctional proteins—such as TJP1, CLDN5, OCLN, and VE-cadherin (CDH5)—are stably expressed throughout life, CLDN11 exhibits a significant positive correlation with age (Pearson correlation p-value < 0.001, Supplemental Fig. 1). CLDN11, which colocalizes with CLDN5, has been shown to increase permeability of the human brain endothelial cell monolayers when its expression is knocked down [30]. Additionally, integrin β1 (ITGB1), the most abundant integrin in our BMV proteome, increases its expression through maturation, reaching its highest levels in adulthood before declining with age. ITGB1 promotes endothelial sprouting while negatively regulating cell proliferation in developmental stage. In maturing vessels, integrin β1 is essential for the proper localization of VE-cadherin, thereby maintaining cell–cell junction integrity[77]. The increased expression of ITGB1 in adults support the requirement for postnatal vascularization during early developmental stage and the necessity of maintaining BBB integrity in later stages. Further research is needed to understand how different collagen families and age-dependent changes in junctional proteins and integrins affect BBB permeability.

### Transporters important for CNS nutrient disposition and brain development are mainly enriched during early development

Our study represents a pioneering effort to map the development and aging trajectory of nutrient transporters on the human BBB. Previous studies in rodents have shown that amino acids such as l-arginine and l-serine accumulate at higher concentrations in younger brains and that the protein expression of Slc7a1, a l-arginine transporter, and Slc38a5 a glutamine, l-serine transporter are significantly higher at earlier developmental stages [78, 79]. Our present study in humans further supports this early need for high concentration of nutrients in the brain. We found that SLC7A1 and SLC38A5 (SNAT5), were highly expressed in early childhood which also aligns with clinical findings that these amino acids present higher CSF to plasma concentration ratios in infancy relative to later in childhood [80]. In comparison to the previous global proteomic study of the human BBB (Al-Majdoub et al. 2019), this study utilizes samples from early development, enabling us to identify several well-known important nutrient SLC transporters at earlier ages. These include riboflavin transporter SLC52A2 and heme/choline transporter SLC49A2, both of which are enriched in the early developmental stage, as well as SLC5A6, SLC19A3, and SLC52A3, which have not been reported in previous untargeted proteomic studies of the human BBB (Al-Majdoub et al. 2019). Mutations in these transporters are associated with Mendelian Diseases which exhibit severe brain developmental phenotypes. The high expression of these nutrient transporters in development is consistent with the brain’s heightened metabolic needs during rapid growth phases and highlights the critical importance of precise regulation of the expression of these transporters in the BBB for normal brain development.

### ADME transporters on BBB exhibit age-dependent patterns

Many known clinically important ADME transporters, including ABCB1, ABCG2, ABCC4, SLCO2B1, SLCO1A2, and SLC29A1, are expressed in our BMV proteome [54]. In alignment with prior research, ABCB1 and ABCG2 were the most abundant ABC transporters identified. While no significant age-related differences in BBB expression levels were noted in these transporters, our data revealed considerable interindividual variability in expression levels, ranging from 1.45 fmol/µg to 13.78 fmol/µg for ABCB1, and from 0.78 fmol/µg to 8.97 fmol/µg for ABCG2. This large variation could potentially lead to significant variation in the brain exposure and associated effects and toxicities of various xenobiotics and drugs among individuals. Age-dependent variation in the expression was observed for some ADME transporters such as SLCO1A2, ABCC4, and SLC22A6. Despite the controversial results from clinical studies on the benefit of statins in autism or AD, their ability to affect brain cholesterol levels cannot be ignored [81–83]. Results from our study point out the possibility that statins may cross the BBB through SLCO1A2, which has lower expression in the elderly population. Expression changes in BBB ADME transporters may also contribute to neurotoxicity [61]. Studies show that in comparison to younger children, older children tend to be more sensitive to methotrexate-induced neurotoxicity [61]. This may be a result of age-related changes in the expression levels of ABCC4, which plays a role in the efflux of MTX (Fig. 4A). Our proteome analysis also identified species differences in the expression levels of drug transporters between humans and rodents, a critical finding since rodent models extensively used in pre-clinical drug development [84]. This finding underscores the critical need for human-based BBB studies, given that most CNS drug development failures stem from a lack of efficacy, and highlights the gap between preclinical models and human studies which needs to be bridged.

### Age is a significant risk factor for many neurological disorders including AD

In our exploratory research using BMV samples from elderly patients, we discovered that 8.6% (595) of proteins change with aging. Notably, proteins involved in inflammation (leukocyte migration) show a marked increase with age, while zinc transport proteins exhibit a significant decline. Zinc deficiency has been linked to an increased risk of AD, possibly due to heightened inflammation. Our findings reveal a reduction in the expression of zinc transporters SLC30A1, SLC30A5, and SLC30A10 among the elderly. Additionally, we observed that APOE, known for its crucial role in lipid metabolism and its contribution to AD pathology, is highly expressed in BMVs. Its expression is comparable to the BMV marker SLC2A1 (GLUT1) and increases with age. Overexpression of APOE in pericytes can lead to their degeneration, which severely impacts BBB function and results in cognitive decline [85]. However, the implications of elevated APOE expression in BMVs remain controversial, with studies indicating it may induce inflammation and subsequently impair BBB integrity [86]. Our study revealed, for the first time, that VEGFB is absent in early childhood and significantly increases with age, showing even higher levels in AD patients. This finding is consistent with BBB disruption in AD, since VEGFB is essential for BBB repair and its expression may have increased in response to BBB damage. Among the 22 AD GWAS genes expressed in our dataset, over half showed a correlation with age, either positive or negative. These results highlight the dynamic nature of gene expression on the BBB related to aging and AD and offer potential new insights into the molecular mechanisms driving AD progression.

The major limitations of this study include the following. First, there is contamination of other brain cell types in our BMV samples. We did observe some astrocyte proteins in our dataset, likely stemming from astrocytic endfoot projections that make contact with BMVs. This type of potential contamination has been reported before when characterizing the BBB (Supplemental Table 5 ([14, 15]). Nonetheless, we observed clear enrichment of BMVs related genes in our samples when comparing between the BMVs and whole brain via qPCR (Supplemental Fig. 6), suggesting that our isolation method effectively enriched for BMVs while maintaining the distinct cellular profiles. Previous studies have been done to assess transporter proteins at the BBB; however, they’ve been limited to adult brains [13–15]. Our results are consistent with all other studies in demonstrating that SLC2A1, which is a marker of the BBB, is one of the most highly expressed SLCs and ABCB1 and ABCG2 are the two most highly expressed ABC transporters on BBB. However, our BMV proteome analysis also revealed values inconsistent with those found in other studies. For instance, the expression level of SLC7A5 in our dataset was significantly higher (3.05 fmol/µg) compared to the findings from Uchida et al. (2011), which used a targeted proteomic approach and reported an average SLC7A5 expression level of 0.431 fmol/µg. This discrepancy aligns with observations made by Al-Majdoub et al. (2019), who noted differences between targeted and global analysis of SLC7A5 expression levels due to the overestimation caused by signal overlap in label-free experiments. Moreover, we observed no expression of SLC51A, a contrast to its detectability by Al-Majdoub et al. (2019) at 0.45 fmol/µg. This re-emphasizes the challenge in comparing quantitative data across studies. An additional limitation of our study is related to the source of the brain samples, which were obtained from the University of Maryland NeuroBioBank. Variable time after death may pass before tissues are frozen[87], which may result in variable protein degradation in the samples. Finally, in the Elderly group, samples were from a small number of donors, especially for the samples from AD patients (N=3). Studies with larger sample sizes are needed to confirm all of these results.

In conclusion, our study inclusively identified proteins in BMVs which display age-dependent patterns. These findings not only shed light on the physiological transformations that occur in the human brain across development and aging but also suggest potential mechanisms for interindividual variation and age-related changes in cellular metabolism and in drug distribution in the brain as well as sensitivity to the CNS effects of drugs. Through a proof-of-concept study utilizing a published phenytoin PBPK model, we illustrate how drug exposure in the brain can be influenced by differences in transporter expression and BBB permeability. These studies can be expanded to other drugs to enhance predictions regarding CNS drug distribution and optimal dosing in pediatric and geriatric patients, especially for drugs intended as medical countermeasures for children and the elderly, vulnerable populations for whom research poses ethical and logistical challenges. Ultimately, this research highlights the critical need of accounting for age-related changes in drug development and personalized medicine.

## Methods

### Human brain tissue samples

Post-mortem frozen brain cortical tissue samples from thirty-four healthy donors and four Alzheimer’s Disease patients were obtained from the National Institutes of Health NeuroBioBank at the University of Maryland, Baltimore, MD. Tissues were stored at −80°C until the day of microvessel isolation. The demographic information of donors is reported in Supplemental Table 1. The pediatric population was separated into different age groups according to the International Council for Harmonisation guidelines: newborns (PNA 0–28 days, GA>37 weeks), infants (1–24 months old), children (2–12 years old), and adolescents (12–16 years old). Donors over 16 years old were defined as adults and donors over 60 years old were defined as Elderly group. However, due to small sample sizes, the newborn and infant were combined into “Development group” and children, adolescents, and adult were combined into “Adult Group” to increase the power of the study after confirming no proteins were differentially expressed between these groups, respectively.

### Isolation of human brain microvessels

The process of isolating brain microvessels (BMVs) from human brain cortical tissue was conducted with some modifications based on a previously described protocol [88]. The procedure started with using less than 1 gram of tissue which was thawed and homogenized in protease inhibitors (cOmplete protease inhibitor cocktail, Sigma-Aldrich, St. Louis, MO) contained HBSS solution with 20 up- and-down strokes in a Potter-Elvehjem glass homogenizer. The resulting homogenate was centrifuged at 1200 g for 10 min at 4C, and the BMV enriched pellet was resuspended in a 17.5% dextran-70/HBSS solution and centrifuged and centrifuged at 4300 g for 15 min at 4C in a swinging bucket rotor. The myelin-rich layer on the top and the supernatant were aspirated and the pellet was resuspended in HBSS with 1% Bovine Serum Albumin (BSA). This solution was filtered with a 40 μm nylon mesh strainer and the BMVs captured on the strainer were washed with 35 ml of 1% BSA/ HBSS buffer. The BMVs were then collected off the filter with 1% BSA/HBSS buffer and centrifuged at 3000 g for 5 min at 4C. The supernatant was removed and the resulting BMV pellet was immediately frozen at -80°C for further analysis. All the steps were carried out at 4°C or on ice.

### Global proteomics using liquid chromatography tandem mass spectrometry (LC/MS)

The process of LC/MS is based on a previously described protocol [88]. BMV samples were lysed in a 100 mM Tris-HCl buffer (pH 7.8) containing 50 mM dithiothreitol and 2% sodium dodecyl sulfate, followed by a 5-minute incubation at 95°C. Subsequently, the samples underwent sonication using a Branson-rod-typesonicator and were then centrifuged at 14,000 g for 10 minutes. To determine protein concentration, a tryptophan fluorescence assay was employed. For multi-enzyme digestion, 100 μg of protein was utilized in the multi-enzyme digestion filter-aided sample preparation (MED-FASP) protocol, where sequential digestion with LysC and trypsin occurred. Following digestion, the resulting peptides were concentrated via a GeneVac EX-2plus system. Subsequently, peptide separation was achieved using an Ultimate 3000 RSLCnano system, employing an easy spray C18 reversed-phase column (50 cm, ID 75 μm) with a gradient of water/acetonitrile containing 0.1% formic acid over a 145-minute period. The eluted peptides were analyzed with a Top15 method, involving full MS followed by ddMS2 scans, using an Orbitrap Q Exactive HF mass spectrometer (ThermoFisher, Waltham, MA). Data analysis was conducted using MaxQuant version 1.6.10.43, with the complete human proteome extracted from UniProt (September 2020). A false discovery rate of 0.01 was set, and match-between-runs was enabled for enhanced accuracy. Protein abundance quantification was performed using the total protein approach (TPA) for proteins with three or more razor+unique peptides. The mass spectrometry proteomics data have been deposited in the ProteomeXchange Consortium through the PRIDE partner repository.

### Differential expression analysis

Unpairwise differentially expressed proteins were identified using Student’s t-test, followed by Benjamini-Hochberg (BH) FDR correction (Significant as Absolute Log2 Fold Change > Log2(1.5), P-value < 0.1). Differential expression was presented in volcano plots, which were generated with the “EnhancedVolcano” version 1.14.0 package in R[89].

### Weighted gene correlation network analysis Discovery brain proteome

Weighted protein coexpression network of the discovery BMV dataset was derived from the protein abundance values using the blockwiseModules WGCNA function ([35]WGCNA 1.72.5 R package) with the following settings: soft threshold power beta = 6, deepSplit=2, minimum module size = 30, merge cut height = 0.25, TOMDenominator = “min”, a signed network with partitioning about medioids (PAM) respecting the dendrogram. This approach calculates pairwise biweight mid-correlations (bicor, a robust correlation metric) between each protein pair and transforms this correlation matrix into a signed adjacency matrix. After blockwiseModules network construction, 30 modules consisting of 34 or more proteins were detected. Module eigenproteins, which are defined as the first principal component (PC) of each module protein expression, were correlated with age(days) using bicor analysis.

### Gene ontology enrichment analysis

The enriched GO terms (biological process, cellular component, and molecular function) of the differential expressed proteins during the developmental stage and aging process were identified and visualized based on the clusterProfiler version 4.4.4 package in R software using the default value[90].

### Physiologically based pharmacokinetic (PBPK) modeling

Systemic and central nervous system (CNS) drug concentrations were simulated using both whole-body and four-compartment permeability-limited brain models within the Simcyp PBPK Simulator (version 21.1 and 22, Certara, Princeton, NJ, USA). The pharmacological and physiological parameters for the model were adapted from a previously published PBPK model for phenytoin [64]. Simulations were conducted over 10 trials, each consisting of 10 virtual healthy volunteers, using the same demographic and dosing information as the referenced phenytoin model. Default settings for BBB P-glycoprotein (P-gp) abundance (3.25 pmol/mg of brain capillary protein) and permeability (46.44 L/h) were applied to generate baseline curves. To mirror the proteomic data observed in our study, adjustments were made to these parameters, and the results of these modifications are presented with the respective parameter values in Figure 4B-E.

## Supporting information

Supplemental Tables link to Figure 3

## Acknowledgments

We are grateful for you for taking the time to read our manuscript. We are also grateful to Dr Qi Liu, Dr Shiew-Mei Huang, Dr Andrew Yang for helpful feedback and discussion as we developed and conducted this project. Human tissue was obtained from the National Institutes of Health NeuroBioBank at the University of Maryland, Baltimore, MD. Part of this work was previously presented at the Annual Meeting of the American Society for Clinical Pharmacology & Therapeutics (ASCPT) held March 2023, Atlanta, USA. This work was supported, in part, by the National Institute of Health grants UC2HD113474, by the US Food and Drug Administration (FDA) through Medical Countermeasures Initiative, grant U01FD004979, which supports the UCSF-Stanford Center of Excellence in Regulatory Science and Innovation. Certara UK Limited (Simcyp Division) granted access to the Simcyp Simulators through a sponsored academic license. X.Z. was supported, in part, by the appointments to the Research Participation Program at the Center for Drug Evaluation and Research, administered by the Oak Ridge Institute for Science and Education through an interagency agreement between the US Department of Energy and the FDA. A.R. is supported by the National Institute of General Medical Sciences of the National Institutes of Health (NIH) under award number 5T32GM007546.

## Author contributions

X.Z, M.A., N.H and A.R wrote the manuscript. X.Z, M.A., N.H, A.R, B.V, S.W.Y, P.A, K.M.G designed the research. X.Z, M.A. and N.H performed research. X.Z, M.A., N.H, A.R, B.V, E.C and S.W.Y, analyzed the data.

## Declaration of generative AI and AI-assisted technologies

During the preparation of this work, the author(s) used ChatGPT to revise the English writing. After using this tool or service, the author(s) reviewed and edited the content as needed and take(s) full responsibility for the content of the publication.

## References

1. Thomsen, M.S., L.J. Routhe, and T. Moos, The vascular basement membrane in the healthy and pathological brain. Journal of Cerebral Blood Flow and Metabolism, 2017. 37(10): p. 3300–3317.

2. Kadry, H., B. Noorani, and L. Cucullo, A blood-brain barrier overview on structure, function, impairment, and biomarkers of integrity. Fluids and Barriers of the Cns, 2020. 17(1).

3. Luissint, A.C., et al., Tight junctions at the blood brain barrier: physiological architecture and disease-associated dysregulation. Fluids Barriers CNS, 2012. 9(1): p. 23.

4. Knox, E.G., et al., The blood-brain barrier in aging and neurodegeneration. Mol Psychiatry, 2022. 27(6): p. 2659–2673.

5. Morris, M.E., V. Rodriguez-Cruz, and M.A. Felmlee, SLC and ABC Transporters: Expression, Localization, and Species Differences at the Blood-Brain and the Blood-Cerebrospinal Fluid Barriers. AAPS J, 2017. 19(5): p. 1317–1331.

6. Nguyen, Y.T.K., et al., The role of SLC transporters for brain health and disease. Cell Mol Life Sci, 2021. 79(1): p. 20.

7. Khan, N.U., et al., Carrier-mediated transportation through BBB, in Brain targeted drug delivery system. 2019, Elsevier. p. 129–158.

8 Cater, R.J. et al. Structural and molecular basis of choline uptake into the brain by FLVCR2. bioRxiv 2023.

9. Geier, E.G., et al., Structure-based ligand discovery for the Large-neutral Amino Acid Transporter 1, LAT-1. Proc Natl Acad Sci U S A, 2013. 110(14): p. 5480-5.

10. Pardridge, W.M., R.J. Boado, and C.R. Farrell, Brain-type glucose transporter (GLUT-1) is selectively localized to the blood-brain barrier. Studies with quantitative western blotting and in situ hybridization. J Biol Chem, 1990. 265(29): p. 18035–40.

11. Tiani, K.A., P.J. Stover, and M.S. Field, The Role of Brain Barriers in Maintaining Brain Vitamin Levels. Annu Rev Nutr, 2019. 39: p. 147–173.

12. Online Mendelian Inheritance in Man (OMIM®), McKusick-Nathans Institute of Genetic Medicine, Johns Hopkins University (Baltimore, MD), 06/2024. Available from: https://omim.org/.

13. Uchida, Y., et al., Quantitative targeted absolute proteomics of human blood-brain barrier transporters and receptors. Journal of Neurochemistry, 2011. 117(2): p. 333–345.

14. Shawahna, R., et al., Transcriptomic and Quantitative Proteomic Analysis of Transporters and Drug Metabolizing Enzymes in Freshly Isolated Human Brain Microvessels. Molecular Pharmaceutics, 2011. 8(4): p. 1332–1341.

15. Al-Majdoub, Z.M., et al., Proteomic Quantification of Human Blood-Brain Barrier SLC and ABC Transporters in Healthy Individuals and Dementia Patients. Molecular Pharmaceutics, 2019. 16(3): p. 1220–1233.

16. Saunders, N.R., et al., Physiology and molecular biology of barrier mechanisms in the fetal and neonatal brain. J Physiol, 2018. 596(23): p. 5723–5756.

17. Mollgard, K., et al., Brain barriers and functional interfaces with sequential appearance of ABC efflux transporters during human development. Sci Rep, 2017. 7(1): p. 11603.

18. Crouch, E.E., et al., Ensembles of endothelial and mural cells promote angiogenesis in prenatal human brain. Cell, 2022. 185(20): p. 3753–3769 e18.

19. Yang, A.C., et al., A human brain vascular atlas reveals diverse mediators of Alzheimer’s risk. Nature, 2022. 603(7903): p. 885–892.

20. Swain, T.R., Clinical trials for children: some concerns. Indian J Pharmacol, 2014. 46(2): p. 145–6.

21. Ek, C.J., et al., Barriers in the developing brain and Neurotoxicology. Neurotoxicology, 2012. 33(3): p. 586–604.

22. Cheung, K.W.K., et al., A Comprehensive Analysis of Ontogeny of Renal Drug Transporters: mRNA Analyses, Quantitative Proteomics, and Localization. Clin Pharmacol Ther, 2019. 106(5): p. 1083–1092.

23. Stephenson, J., et al., Inflammation in CNS neurodegenerative diseases. Immunology, 2018. 154(2): p. 204–219.

24. Sweeney, M.D., A.P. Sagare, and B.V. Zlokovic, Blood-brain barrier breakdown in Alzheimer disease and other neurodegenerative disorders. Nature Reviews Neurology, 2018. 14(3): p. 133–150.

25. Prakash, R. and S.T. Carmichael, Blood-brain barrier breakdown and neovascularization processes after stroke and traumatic brain injury. Curr Opin Neurol, 2015. 28(6): p. 556–64.

26. Kortekaas, R., et al., Blood-brain barrier dysfunction in parkinsonian midbrain in vivo. Ann Neurol, 2005. 57(2): p. 176–9.

27. Chiu, C., et al., P-glycoprotein expression and amyloid accumulation in human aging and Alzheimer’s disease: preliminary observations. Neurobiol Aging, 2015. 36(9): p. 2475–82.

28. Erdo, F. and P. Krajcsi, Age-Related Functional and Expressional Changes in Efflux Pathways at the Blood-Brain Barrier. Front Aging Neurosci, 2019. 11: p. 196.

29. Thomas, P.D., et al., PANTHER: Making genome-scale phylogenetics accessible to all. Protein Sci, 2022. 31(1): p. 8–22.

30. Uchida, Y., et al., Involvement of Claudin-11 in Disruption of Blood-Brain, -Spinal Cord, and -Arachnoid Barriers in Multiple Sclerosis. Mol Neurobiol, 2019. 56(3): p. 2039–2056.

31. Lyu, Q., et al., SENCR stabilizes vascular endothelial cell adherens junctions through interaction with CKAP4. Proc Natl Acad Sci U S A, 2019. 116(2): p. 546–555.

32. McCandless, E.E., et al., Pathological expression of CXCL12 at the blood-brain barrier correlates with severity of multiple sclerosis. Am J Pathol, 2008. 172(3): p. 799–808.

33. Storey, H., et al., COL4A3/COL4A4 mutations and features in individuals with autosomal recessive Alport syndrome. J Am Soc Nephrol, 2013. 24(12): p. 1945–54.

34. Bradshaw, A.D., The role of SPARC in extracellular matrix assembly. J Cell Commun Signal, 2009. 3(3-4): p. 239–46.

35. Langfelder, P. and S. Horvath, WGCNA: an R package for weighted correlation network analysis. BMC Bioinformatics, 2008. 9: p. 559.

36. Shannon, P., et al., Cytoscape: a software environment for integrated models of biomolecular interaction networks. Genome Res, 2003. 13(11): p. 2498–504.

37. Knickmeyer, R.C., et al., A structural MRI study of human brain development from birth to 2 years. J Neurosci, 2008. 28(47): p. 12176–82.

38. Cohen Kadosh, K., et al., Nutritional Support of Neurodevelopment and Cognitive Function in Infants and Young Children-An Update and Novel Insights. Nutrients, 2021. 13(1).

39. Zajec, M., et al., Identification of Blood-Brain Barrier-Associated Proteins in the Human Brain. J Proteome Res, 2021. 20(1): p. 531–537.

40. He, W. and G. Wu, Metabolism of Amino Acids in the Brain and Their Roles in Regulating Food Intake. Adv Exp Med Biol, 2020. 1265: p. 167–185.

41. Shamberger, R.J., Autism rates associated with nutrition and the WIC program. J Am Coll Nutr, 2011. 30(5): p. 348–53.

42. Piecuch, A.K., P.H. Skarzynski, and H. Skarzynski, A Case Report of Riboflavin Treatment and Cochlear Implants in a 4-Year-Old Girl with Progressive Hearing Loss and Delayed Speech Development: Brown-Vialetto-Van Laere Syndrome. Am J Case Rep, 2023. 24: p. e940439.

43. Zhou, L., Association of vitamin B2 intake with cognitive performance in older adults: a cross-sectional study. J Transl Med, 2023. 21(1): p. 870.

44. Tao, L., et al., Dietary Intake of Riboflavin and Unsaturated Fatty Acid Can Improve the Multi-Domain Cognitive Function in Middle-Aged and Elderly Populations: A 2-Year Prospective Cohort Study. Front Aging Neurosci, 2019. 11: p. 226.

45. Yao, Y., et al., Identification and comparative functional characterization of a new human riboflavin transporter hRFT3 expressed in the brain. J Nutr, 2010. 140(7): p. 1220–6.

46. Cater, R.J., et al., Structural and molecular basis of choline uptake into the brain by FLVCR2. Nature, 2024. 629(8012): p. 704–709.

47. Cohen, B.M., et al., Decreased brain choline uptake in older adults. An in vivo proton magnetic resonance spectroscopy study. JAMA, 1995. 274(11): p. 902–7.

48. Lin, L., et al., SLC transporters as therapeutic targets: emerging opportunities. Nat Rev Drug Discov, 2015. 14(8): p. 543–60.

49. Unsal, Y. and G. Hayran, Impact of Early Intervention with Triiodothyroacetic Acid on Peripheral and Neurodevelopmental Findings in a Boy with MCT8 Deficiency. J Clin Res Pediatr Endocrinol, 2023.

50. Balachandran, R.C., et al., Brain manganese and the balance between essential roles and neurotoxicity. J Biol Chem, 2020. 295(19): p. 6312–6329.

51. Lechpammer, M., et al., Pathology of inherited manganese transporter deficiency. Ann Neurol, 2014. 75(4): p. 608–12.

52. Boycott, K.M., et al., Autosomal-Recessive Intellectual Disability with Cerebellar Atrophy Syndrome Caused by Mutation of the Manganese and Zinc Transporter Gene SLC39A8. Am J Hum Genet, 2015. 97(6): p. 886–93.

53. International Transporter, C., et al., Membrane transporters in drug development. Nat Rev Drug Discov, 2010. 9(3): p. 215–36.

54. Galetin, A., et al., Membrane transporters in drug development and as determinants of precision medicine. Nat Rev Drug Discov, 2024.

55. Fokina, V.M., et al., Role of Uptake Transporters OAT4, OATP2A1, and OATP1A2 in Human Placental Bio-disposition of Pravastatin. J Pharm Sci, 2022. 111(2): p. 505–516.

56. Ronaldson, P.T., et al., Transport Properties of Statins by Organic Anion Transporting Polypeptide 1A2 and Regulation by Transforming Growth Factor-beta Signaling in Human Endothelial Cells. J Pharmacol Exp Ther, 2021. 376(2): p. 148–160.

57. Ho, R.H., et al., Drug and bile acid transporters in rosuvastatin hepatic uptake: function, expression, and pharmacogenetics. Gastroenterology, 2006. 130(6): p. 1793–806.

58. Kavey, R.W., et al., Effectiveness and Safety of Statin Therapy in Children: A Real-World Clinical Practice Experience. CJC Open, 2020. 2(6): p. 473–482.

59. Iwaki, M., et al., Inhibition of Methotrexate Uptake via Organic Anion Transporters OAT1 and OAT3 by Glucuronides of Nonsteroidal Anti-inflammatory Drugs. Biol Pharm Bull, 2017. 40(6): p. 926–931.

60. Sane, R., et al., The effect of ABCG2 and ABCC4 on the pharmacokinetics of methotrexate in the brain. Drug Metab Dispos, 2014. 42(4): p. 537–40.

61. Bhojwani, D., et al., Methotrexate-induced neurotoxicity and leukoencephalopathy in childhood acute lymphoblastic leukemia. J Clin Oncol, 2014. 32(9): p. 949–59.

62. Taylor, O.A., et al., Disparities in Neurotoxicity Risk and Outcomes among Pediatric Acute Lymphoblastic Leukemia Patients. Clin Cancer Res, 2018. 24(20): p. 5012–5017.

63. Mori, S., et al., Rat organic anion transporter 3 (rOAT3) is responsible for brain-to-blood efflux of homovanillic acid at the abluminal membrane of brain capillary endothelial cells. J Cereb Blood Flow Metab, 2003. 23(4): p. 432–40.

64. Gaohua, L., et al., Development of a permeability-limited model of the human brain and cerebrospinal fluid (CSF) to integrate known physiological and biological knowledge: Estimating time varying CSF drug concentrations and their variability using in vitro data. Drug Metab Pharmacokinet, 2016. 31(3): p. 224–33.

65. Iorga, A. and B.Z. Horowitz, Phenytoin Toxicity, in StatPearls. 2024: Treasure Island (FL).

66. Wu, M.F. and W.H. Lim, Phenytoin: A Guide to Therapeutic Drug Monitoring. Proceedings of Singapore Healthcare, 2013. 22(3): p. 198–202.

67. Muffat, J. and D.W. Walker, Apolipoprotein D: an overview of its role in aging and age-related diseases. Cell Cycle, 2010. 9(2): p. 269–73.

68. Schachtschneider, K.M., et al., Altered Hippocampal Epigenetic Regulation Underlying Reduced Cognitive Development in Response to Early Life Environmental Insults. Genes (Basel), 2020. 11(2).

69. Watt, N.T., I.J. Whitehouse, and N.M. Hooper, The role of zinc in Alzheimer’s disease. Int J Alzheimers Dis, 2010. 2011: p. 971021.

70. Alsaqati, M., R.S. Thomas, and E.J. Kidd, Proteins Involved in Endocytosis Are Upregulated by Ageing in the Normal Human Brain: Implications for the Development of Alzheimer’s Disease. J Gerontol A Biol Sci Med Sci, 2018. 73(3): p. 289–298.

71. Hsu, C.Y., et al., WWOX and Its Binding Proteins in Neurodegeneration. Cells, 2021. 10(7).

72. Bertram, L. and R.E. Tanzi, Genome-wide association studies in Alzheimer’s disease. Hum Mol Genet, 2009. 18(R2): p. R137–45.

73. Zhao, H., et al., Destabilizing heterochromatin by APOE mediates senescence. Nat Aging, 2022. 2(4): p. 303–316.

74. Mahoney, E.R., et al., Brain expression of the vascular endothelial growth factor gene family in cognitive aging and alzheimer’s disease. Mol Psychiatry, 2021. 26(3): p. 888–896.

75. Ricard-Blum, S., The collagen family. Cold Spring Harb Perspect Biol, 2011. 3(1): p. a004978.

76. Cescon, M., et al., Collagen VI sustains cell stemness and chemotherapy resistance in glioblastoma. Cell Mol Life Sci, 2023. 80(8): p. 233.

77. Yamamoto, H., et al., Integrin beta1 controls VE-cadherin localization and blood vessel stability. Nat Commun, 2015. 6: p. 6429.

78. Radzishevsky, I., et al., Impairment of serine transport across the blood-brain barrier by deletion of Slc38a5 causes developmental delay and motor dysfunction. Proc Natl Acad Sci U S A, 2023. 120(42): p. e2302780120.

79. Tachikawa, M., et al., Developmental changes of l-arginine transport at the blood-brain barrier in rats. Microvasc Res, 2018. 117: p. 16–21.

80. Scholl-Burgi, S., et al., Amino acid cerebrospinal fluid/plasma ratios in children: influence of age, gender, and antiepileptic medication. Pediatrics, 2008. 121(4): p. e920–6.

81. Avan, R., et al., Update on Statin Treatment in Patients with Neuropsychiatric Disorders. Life (Basel), 2021. 11(12).

82. Torrandell-Haro, G., et al., Statin therapy and risk of Alzheimer’s and age-related neurodegenerative diseases. Alzheimers Dement (N Y), 2020. 6(1): p. e12108.

83. Hussain, H.M., M. Zakria, and A.R. Arshad, Statins, incident Alzheimer disease, change in cognitive function, and neuropathology. Neurology, 2008. 71(24): p. 2019; author reply 2019-20.

84. Yang, A.C., et al., Physiological blood-brain transport is impaired with age by a shift in transcytosis. Nature, 2020. 583(7816): p. 425–430.

85. Blanchard, J.W., et al., Reconstruction of the human blood-brain barrier in vitro reveals a pathogenic mechanism of APOE4 in pericytes. Nat Med, 2020. 26(6): p. 952–963.

86. Rieker, C., et al., Apolipoprotein E4 Expression Causes Gain of Toxic Function in Isogenic Human Induced Pluripotent Stem Cell-Derived Endothelial Cells. Arterioscler Thromb Vasc Biol, 2019. 39(9): p. e195–e207.

87. Maryland, U.o., NeuroBioBank (https://www.medschool.umaryland.edu/btbank/brain-donation/the-tissue-donation-process/).

88. Chen, E.C., et al., High Throughput Screening of a Prescription Drug Library for Inhibitors of Organic Cation Transporter 3, OCT3. Pharm Res, 2022. 39(7): p. 1599–1613.

89. Blighe K, R.S., Lewis M, EnhancedVolcano: Publication-ready volcano plots with enhanced colouring and labeling. 2023.

90. Yu, G., et al., clusterProfiler: an R package for comparing biological themes among gene clusters. OMICS, 2012. 16(5): p. 284–7.

