## Supplemental figures for "Proteomic Profiling Reveals Age-Related Changes in Transporter Proteins in the Human Blood-Brain Barrier"

### Slide 1
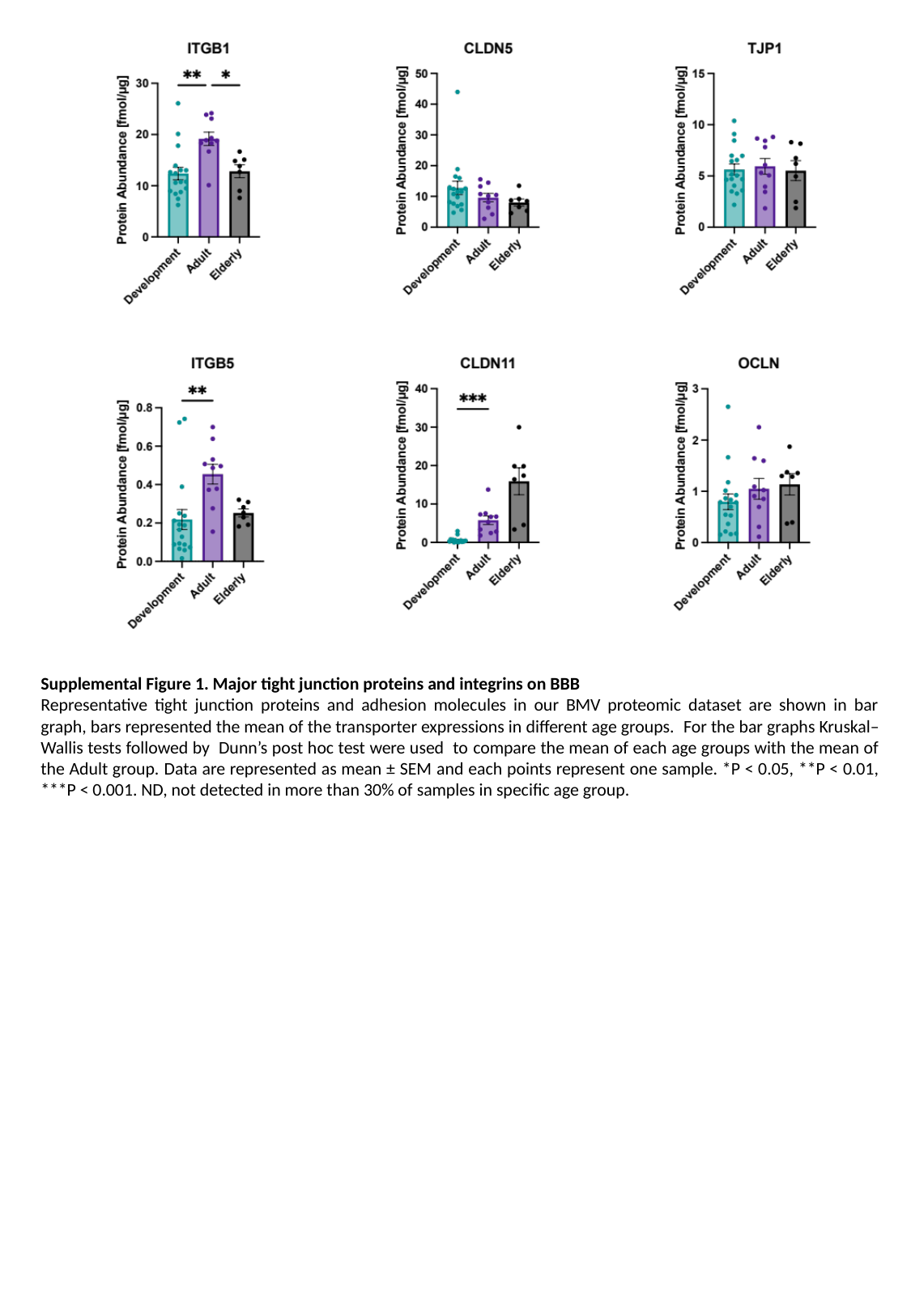

Supplemental Figure 1. Major tight junction proteins and integrins on BBB
Representative tight junction proteins and adhesion molecules in our BMV proteomic dataset are shown in bar graph, bars represented the mean of the transporter expressions in different age groups. For the bar graphs Kruskal–Wallis tests followed by Dunn’s post hoc test were used to compare the mean of each age groups with the mean of the Adult group. Data are represented as mean ± SEM and each points represent one sample. *P < 0.05, **P < 0.01, ***P < 0.001. ND, not detected in more than 30% of samples in specific age group.

### Slide 2
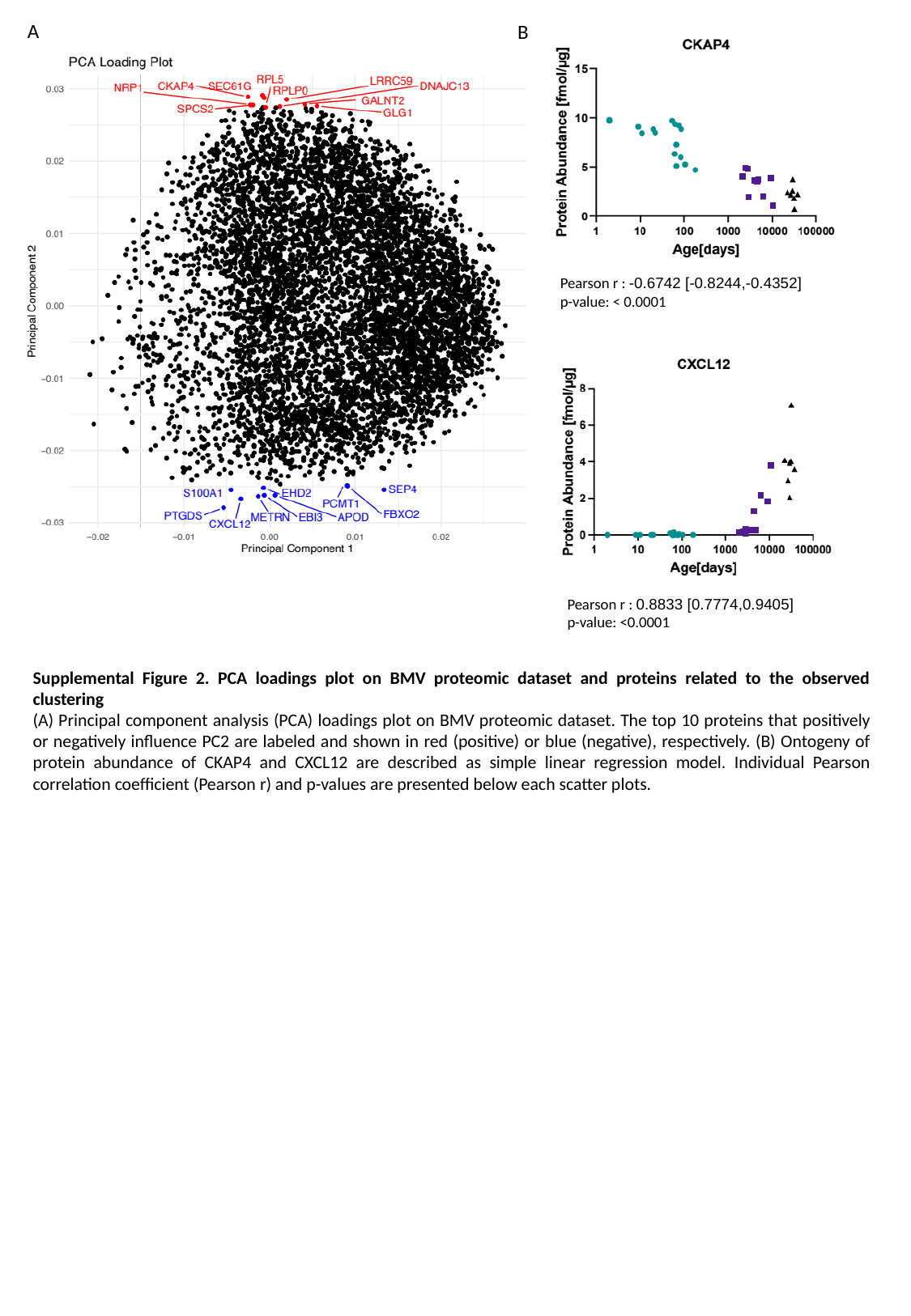

A
B
Pearson r : -0.6742 [-0.8244,-0.4352]
p-value: < 0.0001
Pearson r : 0.8833 [0.7774,0.9405]
p-value: <0.0001
Supplemental Figure 2. PCA loadings plot on BMV proteomic dataset and proteins related to the observed clustering
(A) Principal component analysis (PCA) loadings plot on BMV proteomic dataset. The top 10 proteins that positively or negatively influence PC2 are labeled and shown in red (positive) or blue (negative), respectively. (B) Ontogeny of protein abundance of CKAP4 and CXCL12 are described as simple linear regression model. Individual Pearson correlation coefficient (Pearson r) and p-values are presented below each scatter plots.

### Slide 3
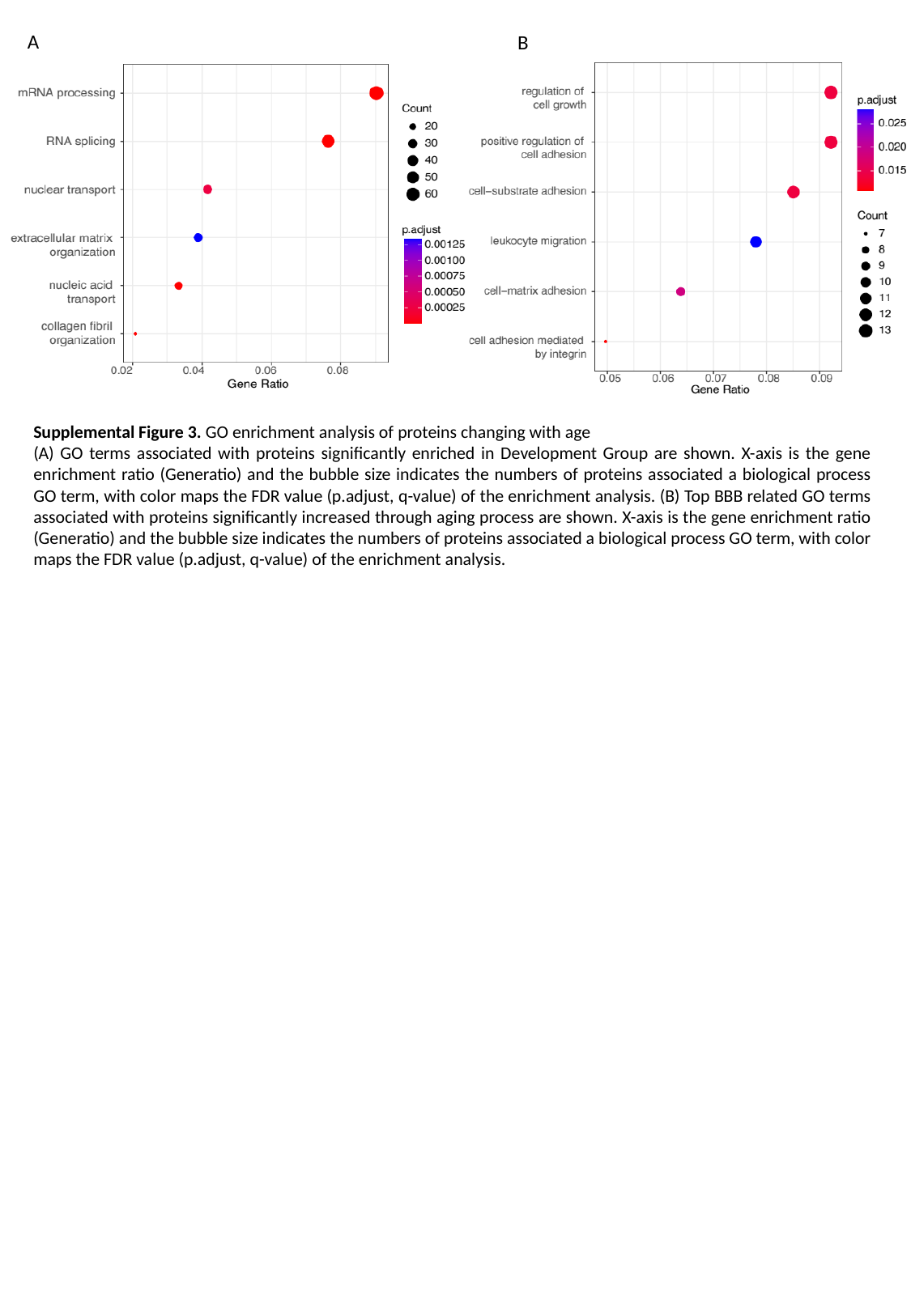

A
B
Supplemental Figure 3. GO enrichment analysis of proteins changing with age
(A) GO terms associated with proteins significantly enriched in Development Group are shown. X-axis is the gene enrichment ratio (Generatio) and the bubble size indicates the numbers of proteins associated a biological process GO term, with color maps the FDR value (p.adjust, q-value) of the enrichment analysis. (B) Top BBB related GO terms associated with proteins significantly increased through aging process are shown. X-axis is the gene enrichment ratio (Generatio) and the bubble size indicates the numbers of proteins associated a biological process GO term, with color maps the FDR value (p.adjust, q-value) of the enrichment analysis.

### Slide 4
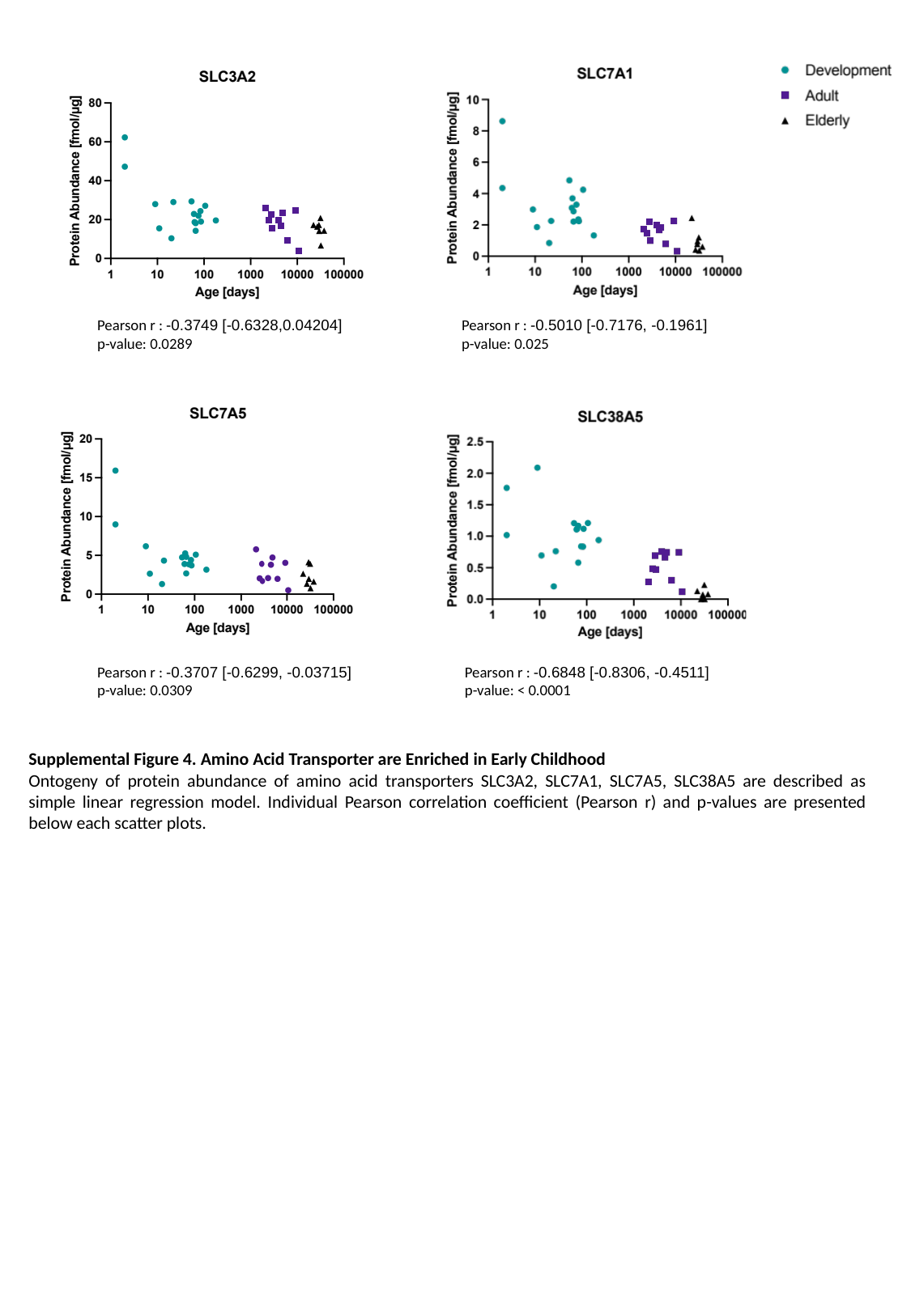

Pearson r : -0.3749 [-0.6328,0.04204]
p-value: 0.0289
Pearson r : -0.5010 [-0.7176, -0.1961]
p-value: 0.025
Pearson r : -0.3707 [-0.6299, -0.03715]
p-value: 0.0309
Pearson r : -0.6848 [-0.8306, -0.4511]
p-value: < 0.0001
Supplemental Figure 4. Amino Acid Transporter are Enriched in Early Childhood
Ontogeny of protein abundance of amino acid transporters SLC3A2, SLC7A1, SLC7A5, SLC38A5 are described as simple linear regression model. Individual Pearson correlation coefficient (Pearson r) and p-values are presented below each scatter plots.

### Slide 5
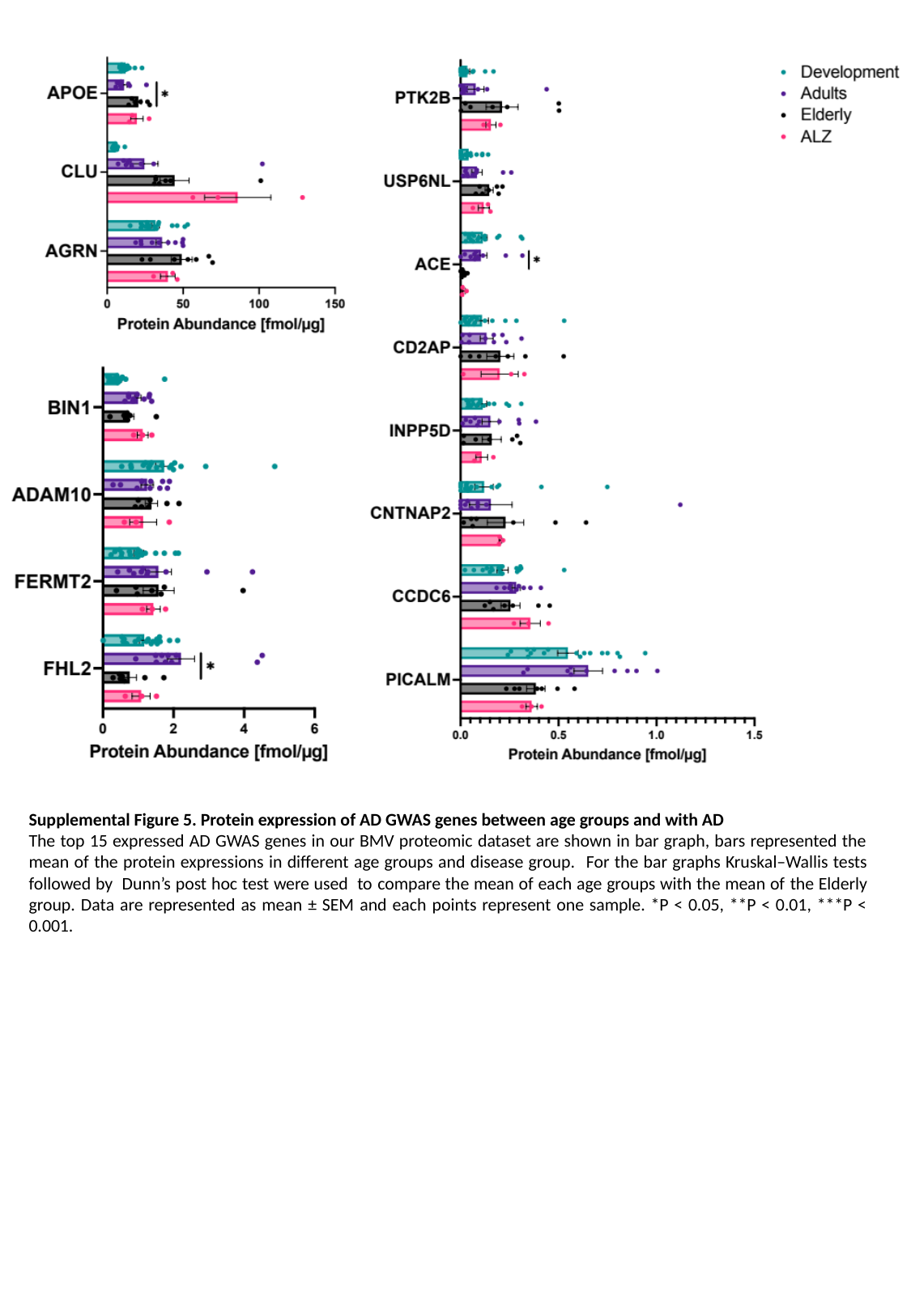

Supplemental Figure 5. Protein expression of AD GWAS genes between age groups and with AD
The top 15 expressed AD GWAS genes in our BMV proteomic dataset are shown in bar graph, bars represented the mean of the protein expressions in different age groups and disease group. For the bar graphs Kruskal–Wallis tests followed by Dunn’s post hoc test were used to compare the mean of each age groups with the mean of the Elderly group. Data are represented as mean ± SEM and each points represent one sample. *P < 0.05, **P < 0.01, ***P < 0.001.

### Slide 6
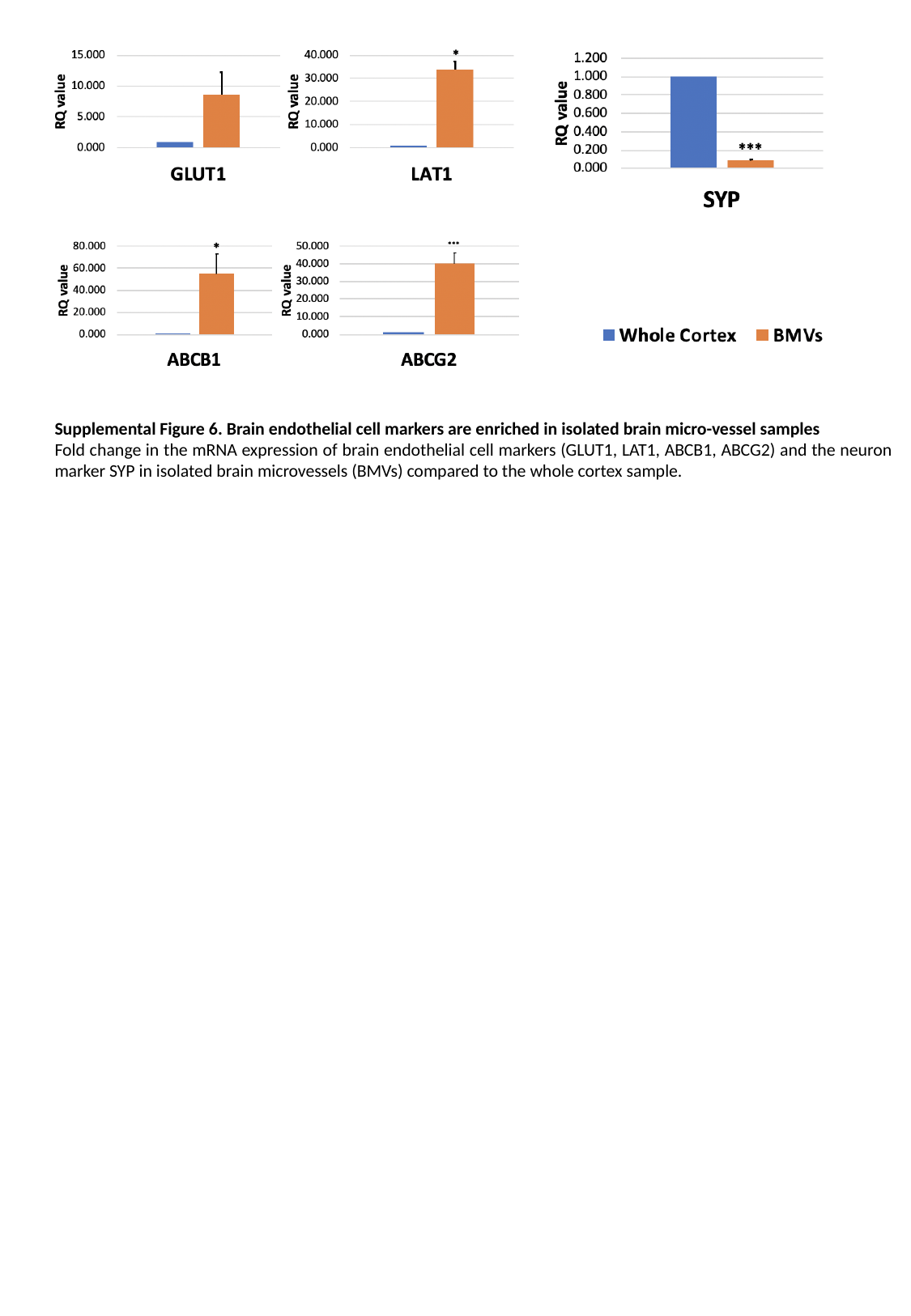

Supplemental Figure 6. Brain endothelial cell markers are enriched in isolated brain micro-vessel samples
Fold change in the mRNA expression of brain endothelial cell markers (GLUT1, LAT1, ABCB1, ABCG2) and the neuron marker SYP in isolated brain microvessels (BMVs) compared to the whole cortex sample.
