## Supplemental Tables for "Proteomic Profiling Reveals Age-Related Changes in Transporter Proteins in the Human Blood-Brain Barrier"

**Supplemental Table 1. Overview of sample size and age range of samples**

| **Age group** |  | **Number of samples** | **Age range** |
| --- | --- | --- | --- |
| Development | Fetus/Neonate | 7 | -37 weeks gestation – 28 days |
|  | Infant | 10 | 1 – 6 months |
| Adult | Child | 5 | 4 – 12 years |
|  | Adolescent/Adult | 5 | 12 – 30 years |
| Elderly | Adult >60yrs | 7 | 61 – 103 years |
| - | Alzheimer’s disease | 3 | 72 – 90 years |
|  | Total | 38 |  |

**Supplemental Table 2. Macronutrients transporters on BBB**

|  |  |  | **Protein Abundance [pmol/mg]** | | |  |  |
| --- | --- | --- | --- | --- | --- | --- | --- |
| **Substrate Category** | **Gene (Protein)** | **Predominant substrate(s)** | **Development** | **Adult** | **Elderly** | **Developmental Changes *(p-value)** | **Aging Changes *(p-value)** |
| **Amino Acids** | SLC1A1(EAAT3) | L-Glu, D/L-Asp, L- Cys | ND | 0.033 | 0.367 | ↑ | 0.160 |
|  | SLC1A2(EAAT2) | L-Glu, D/L-Asp | 9.621 | 11.144 | 12.447 | 0.696 | 0.733 |
|  | SLC1A3(EAAT1) | L-Glu, D/L-Asp | 11.564 | 8.108 | 9.718 | 0.213 | 0.590 |
|  | SLC1A4(ASCT1) | L-Ala, L-Ser | 1.305 | 0.393 | 0.692 | ↓(0.0423) | 0.065 |
|  | SLC1A5(ASCT2) | L-Asp, L-Cys, L-Gln | ND | ND | 0.044 | - | ↑ |
|  | SLC3A2  (CD98hc/4F2hc) | - | 26.093 | 18.151 | 15.261 | ↓(0.048) | 0.313 |
|  | SLC6A17(NTT4) | NAAs | 0.125 | 0.178 | 0.176 | 0.655 | 0.988 |
|  | SLC7A1(CAT1) | CAAs (L-Arg) | 3.318 | 1.527 | 0.970 | ↓(0.0014) | 0.123 |
|  | SLC7A5(LAT1) | LNAAs | 5.235 | 3.054 | 2.346 | ↓(0.032) | 0.333 |
|  | SLC7A8(LAT2) | LNAAs | 0.462 | 0.126 | 0.028 | ↓(0.000011) | ↓(0.0066) |
|  | SLC7A11 (xCT) | L-Glu, L-Cys | 0.037 | 0.027 | 0.018 | 0.231 | 0.276 |
|  | SLC7A14 | CAAs | ND | 0.073 | 0.115 | ↑ | 0.602 |
|  | SLC25A12  (AGC1) | L-Glu, D/L-Asp | 6.281 | 4.464 | 5.359 | 0.078 | 0.528 |
|  | SLC25A13  (AGC2/Aralar2) | L-Asp, L-Glu | 1.393 | 0.745 | 0.983 | ↓(0.00036) | 0.355 |
|  | SLC25A15  (ORNT1) | L-Orn, L-Cit | 0.055 | 0.044 | ND | 0.706 | ↓ |
|  | SLC25A22  (GC-1) | L-Glu | 6.055 | 4.086 | 3.935 | 0.282 | 0.947 |
|  | SLC32A1  (VIAAT) | L-Gly, GABA | 0.251 | 0.250 | 0.190 | 0.997 | 0.793 |
|  | SLC38A1  (SNAT1) | L-Gln | 0.020 | ND | ND | ↓ | - |
|  | SLC38A2  (SNAT2) | L-Gln | 0.090 | 0.082 | 0.019 | 0.749 | ↓(0.0032) |
|  | SLC38A3  (SNAT3) | L-Gln | 1.220 | 0.566 | 0.555 | ↓(0.0015) | 0.943 |
|  | SLC38A5  (SNAT5) | L-Gln, L-Ser | 1.027 | 0.527 | 0.082 | ↓(0.00062) | ↓(0.00012) |
|  | SLC38A10 | L-Gln, L-Ala | ND | 0.020 | ND | ↑ | ↓ |
| **Glucose** | SLC43A2(LAT4) | BCAAs | ND | 0.040 | ND | ↑ | ↓ |
|  | SLC2A1  (GLUT1)) | Glucose | 25.855 | 38.187 | 18.229 | 0.075 | ↓(0.0059) |
|  | SLC2A3(GLUT3) | Glucose | 1.185 | 1.109 | 1.670 | 0.844 | 0.293 |
|  | SLC2A4(GLUT4) | Glucose | 0.012 | 0.125 | 0.104 | ↑(0.017) | 0.643 |
| **Nucleotide sugars** | SLC5A3(SMIT1) | Myoinositol (glucose) | 0.073 | 0.062 | ND | 0.731 | ↓ |
|  | SLC35A4  (MGC2541) | PAPS | 0.357 | 1.137 | 1.604 | ↑(0.0054) | 0.315 |
|  | SLC35B1  (UGTREL1) | UDPGA | 0.041 | 0.019 | ND | ↓(0.017) | ↓ |
|  | SLC35B2  (PAPST1) | PAPS | 0.086 | 0.047 | 0.059 | ↓(0.011) | 0.704 |
| **Choline** | SLC49A2**  (FLVCR2) | Choline, heme | 0.030 | 0.013 | ND | ↓(0.035) | ↓ |
|  | SLC44A1 | Choline | 0.483 | 0.945 | 1.228 | ↑(0.0065) | 0.144 |
|  | SLC44A2 | Choline | 2.072 | 2.133 | 2.002 | 0.810 | 0.380 |
| **Others** | SLC59A1  (MFSD2A) | DHA | 0.0672 | ND | ND | ↓ | - |

*P-values are calculated with Welch’s t-test. ** Detected with less three than razor + unique peptides. Abbreviations: BCAA, Branched-chain amino acids; CAA, cationic amino acid; DHA, docosahexaenoic acid; LCFA, Long-chain fatty acids; NAA, neutral amino acid; PAPS, 3'-phosphoadenosine 5'-phosphosulfate; UDPGA, uridine 5′-diphospho-α-D-glucuronic acid. ND, not detected in more than 30% of samples in specific age group.

**Supplemental Table 3. Micronutrient transporters on BBB**

|  |  |  | **Protein Abundance [pmol/mg]** | | |  |  |
| --- | --- | --- | --- | --- | --- | --- | --- |
| **Substrate Category** | **Gene (Protein)** | **Predominant substrate(s)** | **Development** | **Adult** | **Elderly** | **Developmental Changes *(p-value)** | **Aging Changes *(p-value)** |
| **Vitamins** | SLC19A1  (RFC) | Folates | 0.199 | 0.495 | 0.331 | ↑(0.012) | 0.195 |
|  | SLC19A3  (THTR2) | Thiamine | ND | 0.218 | 0.167 | ↑ | 0.480 |
|  | SLC25A19  (DNC) | Thiamine pyrophosphate | 0.021 | 0.019 | 0.006 | 0.767 | 0.081 |
|  | SLC52A2**  (RFVT2) | Riboflavin | 0.340 | 0.169 | ND | ↓(0.02) | ↓ |
|  | SLC52A3  (RFVT3) | Riboflavin | ND | ND | 0.245 | - | ↑ |
| **Metals** | SLC30A1  (ZNT1) | Zinc | 1.255 | 1.279 | 0.642 | 0.926 | ↓(0.007) |
|  | SLC30A3  (ZNT3) | Zinc | 0.331 | 0.529 | ND | 0.384 | ↓ |
|  | SLC30A5**  (ZNT5) | Zinc | 0.085 | 0.045 | ND | ↓(0.007) | ↓ |
|  | SLC30A7  (ZNT7) | Zinc | 0.308 | 0.126 | 0.114 | ↓(0.002) | 0.809 |
|  | SLC30A9  (ZNT9) | Zinc | 0.185 | 0.221 | 0.127 | 0.449 | 0.150 |
|  | SLC30A10  (ZNT10) | Manganese, Zinc | 0.167 | 0.078 | ND | ↓(0.012) | ↓ |
|  | SLC39A6  (ZIP6) | Zinc | 0.028 | ND | ND | ↓ | - |
|  | SLC39A7**  (ZIP7) | Zinc, Manganese | 0.274 | 0.167 | 0.199 | 0.154 | 0.672 |
|  | SLC39A8**  (ZIP8) | Zinc, Cadmium, Manganese | 0.053 | 0.094 | ND | 0.057 | ↓ |
|  | SLC39A10  (ZIP10) | Zinc | 0.116 | 0.129 | 0.019 | 0.640 | ↓(<0.001) |
|  | SLC39A11** (ZIP11) | Zinc | ND | ND | 0.202 | - | ↑ |
|  | SLC39A14** (ZIP14) | Zinc, Iron, Cadmium, Manganese | 0.124 | 0.057 | 0.128 | 0.067 | 0.054 |
|  | SLC40A1  (FPN1) | Ferrous iron | 0.104 | 0.078 | ND | 0.396 | ↓ |
|  | TFRC (TFR1) | Iron | 13.382 | 5.915 | 2.870 | ↓(<0.001) | ↓(0.005) |
| **Neuro-**  **transmmiters** | SLC6A1  (GAT1) | GABA | 1.854 | 2.972 | 1.630 | ↓(0.021) | ↓(0.026) |
|  | SLC6A12  (BGT1) | betaine, GABA | 1.473 | 2.506 | 2.409 | ↓(0.018) | 0.883 |
|  | SLC6A13  (GAT2) | betaine, GABA | 0.533 | 0.390 | 0.733 | 0.238 | 0.233 |

*P-values are calculated with Welch’s t-test. ** Detected with less than three razor + unique peptides. Abbreviations: GABA, Gamma-aminobutyric acid. ND, not detected in more than 30% of samples in specific age group.

**Supplemental Table 4. Non-detected ADME transporters in our BMV proteome**

| **Family** | **Non-detected in our dataset** |
| --- | --- |
| MRP | ABCC1, ABCC2, ABCC3 |
| OCT | SLC22A1(OCT1), SLC22A2(OCT3) |
| OAT | SLC22A7(OAT2), SLC22A8(OAT3), SLC22A9(OAT4) |
| OCTN | SLC22A5(OCTN2) |
| MDR | ABCB4 |
| PEPT | SLC15A1, SLC15A2 |
| MATE | SLC47A1, SLC47A2 |
| OST | OST α/β |
| OATP | SLCO1B1, SLCO1B3, SLC4C1 |

Abbreviations: MATE, Multidrug and toxin extrusion; MDR, Multidrug resistance; MRP, Multidrug resistance-associated protein; OAT, Organic anion transporter; OATP, Organic anion transporting polypeptide; OCT, Organic cation transporter; OCTN, Organic cation/carnitine transporter; OST, Organic solute transporter; PEPT, Peptide transporter.

**Supplemental Table 5. Compare to previous studies (Markers for other cell types)**

| **Markers** | **Cell type** | **The present study [fmol/µg]** | **Al-Majdoub et al,2019 [fmol/µg]** | **Shawhana et al., 2011 [fmol/µg]** | **Uchida et al., 2011 [fmol/µg]** |
| --- | --- | --- | --- | --- | --- |
| **GFAP** | Astrocytes | 205.22 ± 213.77 | 19.2 ± 18.4 | 503.2 ± 174.09 | NQ |
| **NG2(CSPG4)** | Pericytes | 1.21 ± 0.55 | 0.16 ± 0.06 | 1.07 ± 0.41 | NQ |
| **SYP** | Neurons | 11.21 ± 12.57 | 3.01 ± 2.45 | 1.45 ± 0.37 | NQ |
